# Early-life canine gut microbiome maturation follows a shared age–diet trajectory within persistent host-specific structure

**DOI:** 10.64898/2026.05.25.727648

**Authors:** Tamás Járay, Gábor Gulyás, Md Asaduzzaman, Ákos Dörmő, Zsolt Csabai, Balázs Kakuk, Zsolt Boldogkői, Dóra Tombácz

**Affiliations:** Department of Medical Biology, Albert Szent-Györgyi Medical School, University of Szeged, Somogyi u. 4., 6720 Szeged, Hungary; MTA-SZTE Lendület GeMiNI Research Group, University of Szeged, Somogyi u. 4., 6720 Szeged, Hungary

**Author notes:** **Corresponding author**: DT. **Emails:** TJ GG MA ÁD ZC BK ZB.

**Keywords:** canine gut microbiome, early-life maturation, host identity, shotgun metagenomics, long-read 16S sequencing, virome, functional inference

## Abstract

Early-life gut microbiome maturation remains poorly resolved in dogs. We analyzed 146 fecal samples from 19 Pumi puppies across four dietary stages using paired shotgun metagenomics and Oxford Nanopore full-length 16S sequencing. Bacterial communities shifted from *Escherichia*-rich facultative assemblages during breastfeeding to anaerobe-rich communities after the transition to solid food, with the strongest compositional change occurring during dietary diversification. Functional profiles changed most strongly at the start of complementary feeding, while viral profiles showed increasing phage richness and stage-dependent predicted phage–host associations. Despite this shared dietary trajectory, host identity explained a substantial fraction of microbiome variation and produced persistent dog-specific abundance patterns. Long-read 16S profiles recovered the main developmental signal despite platform-dependent taxonomic bias, and 16S-based functional inference captured pathway-level contrasts concordant with matched metagenomes. These results identify early-life canine gut microbiome maturation as an age-diet-associated but host-constrained ecological process involving coordinated bacterial, viral, and functional restructuring.

## Introduction

Early-life maturation of the gut microbiome shapes immune, metabolic, and developmental trajectories across mammalian hosts. Microbial communities are influenced by genetics, diet, environment, and host–microbe interactions (Singh et al., 2017; Ferretti et al., 2018; Xu et al., 2020; Tavalire et al., 2021). Altered gut microbial structure and function have been associated with metabolic and autoimmune disorders, as well as altered susceptibility to infections (Sanz et al., 2015; Shaheen et al., 2022; Bernard-Raichon et al., 2022). In humans, the gut microbiome is highly dynamic during infancy, with rapid ecological turnover coinciding with immune maturation and metabolic development, potentially influencing long-term health and disease risk (Hill et al., 2017; Liu et al., 2025).

Longitudinal, high-resolution microbiome datasets remain scarce outside human cohorts, particularly in translational animal models. Rodents are widely used but differ substantially from humans in diet, gastrointestinal anatomy, and microbiota composition. In contrast, dogs (*Canis lupus familiaris*) share several physiological, dietary, and environmental features with humans (Coelho et al., 2018; Hernandez et al., 2022), making them a useful model for gut microbiome research (Vázquez-Baeza et al., 2016; Branck et al., 2024). Their shorter lifespan enables dense longitudinal sampling across developmental stages, while controlled breeding programs allow partial standardization of genetic background, housing, and feeding conditions.

Most canine microbiome research has focused on adult dogs or cross-sectional cohorts (Suchodolski, 2011; Moon et al., 2018; Masuoka et al., 2017; You and Kim, 2021; Omatsu et al., 2018; Vilson et al., 2018). Early-life microbial dynamics, including maternal seeding, postnatal colonization, and weaning-associated transitions, remain less well characterized (Pilla and Suchodolski, 2020; Guard et al., 2017). Previous studies have described developmental stages and diet-associated shifts in puppies (Buddington, 2003; Blake et al., 2020; Tal et al., 2021; Pereira et al., 2020), but few longitudinal canine studies have integrated taxonomic, functional, and viral profiles within controlled cohorts spanning breastfeeding, complementary feeding, weaning transition, and established solid diet stages.

Beyond community-wide bacterial and viral dynamics, inter-individual variation represents another important axis of early-life gut ecosystem structuring. Host identity can strongly influence microbial composition, even under shared environmental and dietary conditions. In longitudinal designs, such host effects can appear as individualized abundance baselines layered onto global age- or diet-associated shifts. Although PERMANOVA can quantify the contribution of host identity to overall compositional variance, it does not estimate host-associated variance at the individual taxon level. Mixed-effects modeling addresses this limitation by decomposing variance into between-host and within-host components. By modeling host identity as a random effect, taxon-wise mixed-effects models can quantify host-associated abundance variation after accounting for age or diet. This approach complements community-level analyses and remains rarely applied in canine early-life cohorts; apart from limited reports (e.g., Sweeny et al., 2023), systematic host-effect decomposition in controlled puppy populations is largely lacking.

Interpreting developmental and host-specific microbiome dynamics requires rigorous taxonomic and functional profiling. Metagenomic whole-genome shotgun sequencing (mWGS) is widely regarded as the reference standard because it directly quantifies genes and pathways independent of phylogenetic inference. However, mWGS remains costly and computationally intensive, which can limit scalability in longitudinal designs. Long-read sequencing of full-length V1–V9 16S rRNA genes using Oxford Nanopore Technologies (ONT) provides an alternative with improved taxonomic resolution compared with short-read amplicon sequencing. However, 16S-based approaches remain susceptible to PCR bias and variation in ribosomal operon copy number across taxa, which can distort relative abundance estimates.

Ribosomal operon copy-number normalization using resources such as rrnDB has been proposed to reduce this bias by approximating genome-equivalent abundances. Its effect on full-length ONT 16S data remains unclear, however, and it is not established whether normalization consistently improves concordance with mWGS across taxa. Incomplete database coverage and strain-level variability may introduce additional bias. To date, no systematic evaluation has quantified the effects of rrnDB normalization on taxonomic agreement between ONT V1–V9 profiles and matched mWGS data at both species and community levels.

Functional inference from 16S data presents an additional methodological challenge. Tools such as PICRUSt2 infer gene content from phylogenetic placement, providing a cost-effective alternative to mWGS-based functional profiling. Although PICRUSt2 has been validated mainly on short-read Illumina datasets, its performance with full-length ONT V1–V9 data remains insufficiently characterized. Improved taxonomic resolution could enhance functional inference by enabling more precise trait mapping. However, preprocessing strategies for long-read data, including clustering-based consensus generation and amplicon sequence variant reconstruction, may influence phylogenetic placement and downstream functional predictions.

Simple correlations between predicted and observed functional profiles can overestimate agreement because of compositional constraints and limited variance. The contrast-based framework proposed by Sun et al. (2020) addresses this by testing whether predicted and observed profiles support the same biological differences between groups. Systematic benchmarking of ONT-based functional inference against matched shotgun metagenomes using this type of inference-level validation remains lacking.

Here, we analyzed gut microbiome maturation in a longitudinal cohort of Pumi puppies spanning breastfeeding, complementary feeding, weaning transition, and established solid diet stages. Using paired Illumina shotgun metagenomics and ONT full-length 16S sequencing, we characterized bacterial succession, functional restructuring, virome dynamics, and host-associated microbiome variation across early development. We further benchmarked ONT-derived taxonomic and functional profiles against matched metagenomes to evaluate the extent to which long-read 16S workflows recover biologically relevant developmental signals. This design provides an integrated view of early-life canine gut microbiome maturation and its host-specific structuring.

## Results

### 1. Dietary stage drives shared maturation of the early-life gut microbiome

We generated 146 ONT full-length 16S and 146 Illumina shotgun metagenomic libraries from fecal samples collected from 19 Pumi puppies across four dietary stages. Following quality filtering and downstream preprocessing, 142 matched ONT–mWGS sample pairs were retained for cross-platform analyses. On average, mWGS yielded approximately 1.6 million read pairs per sample, of which approximately 0.78 million were retained after filtering, while ONT sequencing generated approximately 0.4 million full-length reads per sample (**Supplementary Figure S1**; **Supplementary Table S1**).

We first used mWGS-derived taxonomic profiles to define the shared developmental trajectory of the bacterial community across phylum, order, genus, and species levels (**Supplementary Table S2**). Bray–Curtis dissimilarities visualized by NMDS showed progressive separation of samples across dietary stages, with early milk-fed communities most distinct from later solid-diet communities and intermediate stages occupying transitional positions (**Figure 1A**). Hierarchical clustering of pairwise Bray–Curtis distances supported the same stage-associated structure, with the largest between-group dissimilarity observed between Ag1 and Ag4 **(Figure 1B**; **Supplementary Table S3**).

**Figure 1.**
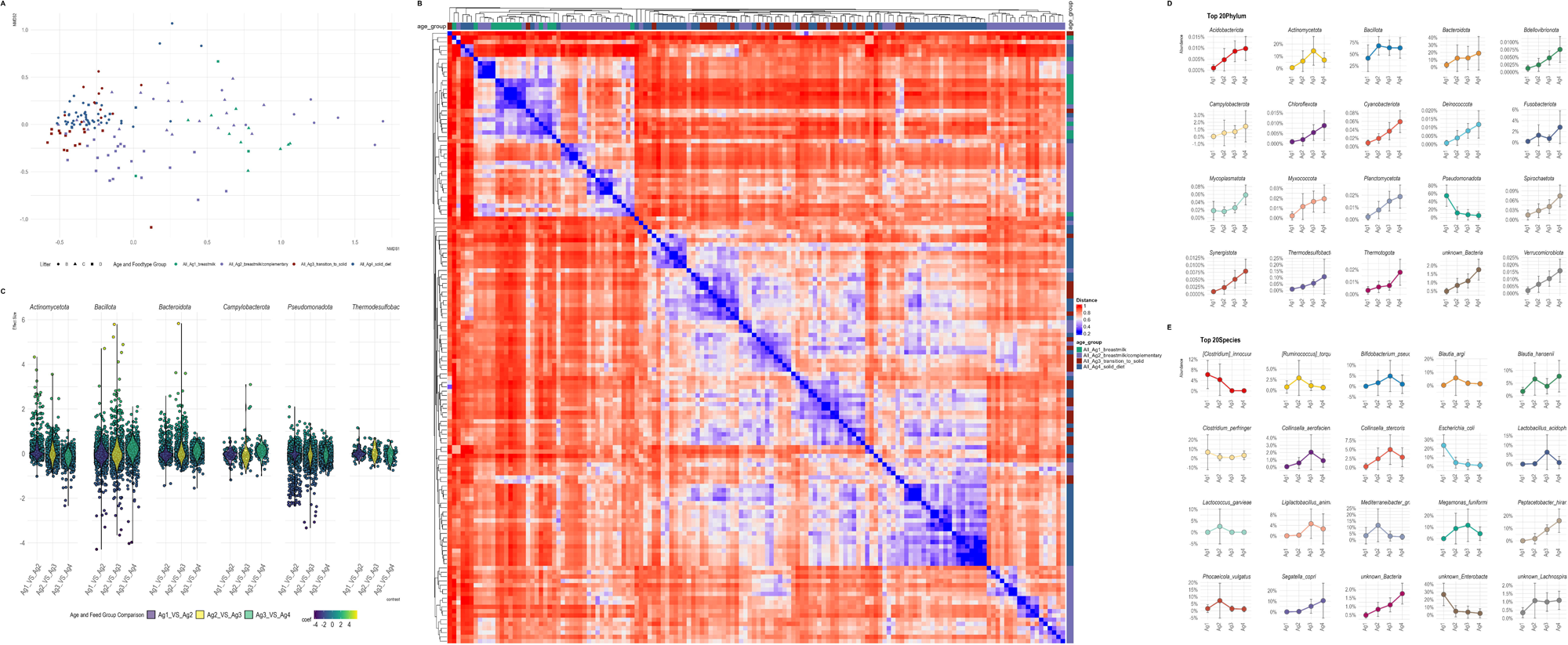
Dietary-stage-associated restructuring of the early-life canine gut microbiome. **A**, NMDS ordination of Bray–Curtis dissimilarities calculated from mWGS-derived taxonomic profiles. Points represent individual samples, colored by developmental stage and shaped by litter. **B**, Hierarchically clustered heatmap of pairwise Bray–Curtis dissimilarities. Color indicates dissimilarity; annotations denote developmental stage. **C**, Distributions of MaAsLin2 effect sizes for bacterial species across consecutive age–diet contrasts, stratified by phylum. Points denote individual species; colors indicate coefficient direction and magnitude. Positive coefficients indicate enrichment in the later stage, whereas negative coefficients indicate enrichment in the earlier stage. **D**, Mean relative abundances of dominant bacterial phyla across developmental stages based on mWGS-derived taxonomic profiles. **E**, Mean relative abundances of the 20 most abundant bacterial species across Ag1–Ag4. Error bars indicate between-sample variability.

Differential abundance analysis with MaAsLin2 identified broad age-associated restructuring across taxonomic ranks. The strongest signal occurred during the Ag2→Ag3 transition, which yielded the highest number of significantly associated taxa at phylum, order, genus, and species levels, consistent with the shift from complementary feeding toward solid diet as the major compositional turning point (**Figure 1C**; **Supplementary Table S4**).

Age-associated restructuring was driven largely by dominant community members. Among the 20 most abundant phyla, approximately 70–75% were significantly associated with at least one developmental transition; at order level, approximately 75–85% of the top 20 orders showed significant age-associated effects (**Figure 1D**; **Supplementary Table S5A–B**).

Species-level profiles provided the clearest biological anchors for this maturation trajectory. Of the 20 most abundant species, 15 were significantly age-associated. *Escherichia coli* declined from 23.5% mean relative abundance in Ag1 to 0.9% in Ag4, whereas anaerobe-associated taxa expanded during later stages. *Segatella copri* increased from 0.35% in Ag2 to 10.5% in Ag4, and *Peptacetobacter hiranonis* increased from 1.7% in Ag2 to 16.0% in Ag4 (**Figure 1E**; **Supplementary Table S5C**). These patterns define a coordinated transition from early facultative assemblages toward an anaerobe-rich, diet-adapted gut microbiome (**Supplementary Results 1**).

### 2. Host identity structures individual maturation trajectories

Although dietary stage defined a shared maturation trajectory, substantial inter-individual variation remained within each stage. We therefore asked whether host identity contributed to microbiome organization beyond the common age–diet effect. PERMANOVA analyses across phylum-, genus-, and species-level profiles showed that dog identity consistently explained a larger fraction of community variation than age/food group, with the host effect becoming most pronounced at finer taxonomic resolution (**Figure 2A**; **Supplementary Table S6**).

**Figure 2.**
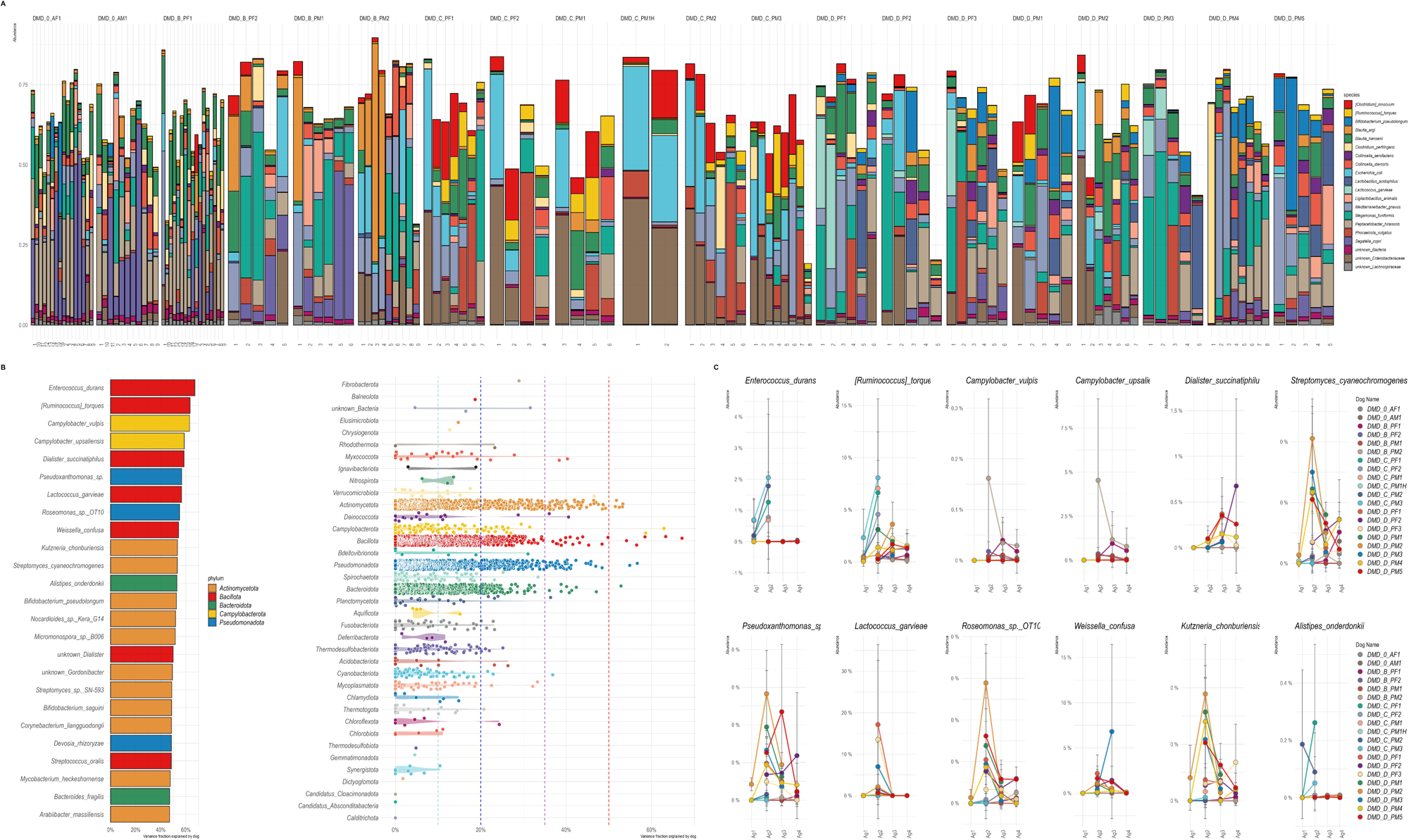
Host identity structures individualized microbiome trajectories. **A**, Species-level relative abundance profiles across individual samples. Stacked bars represent samples grouped by dog; x-axis labels denote sampling order. **B**, Host-associated variance estimated using mixed-effects models. Left, top species ranked by variance explained by dog identity, with bars colored by phylum. Right, distribution of dog-associated variance across taxa at multiple taxonomic levels; each point represents one taxon and dashed lines indicate reference thresholds. **C**, Dog-specific developmental trajectories for selected species across Ag1–Ag4. Each line represents an individual dog; points indicate stage-level means with within-stage variability.

To identify host-associated taxa, we applied taxon-wise CLR-transformed linear mixed-effects models following the conceptual framework of Sweeny et al. (2023). For each taxon, CLR-transformed abundance was modeled as a function of age–diet group, with dog identity included as a random intercept. Host association was summarized as the proportion of model variance attributable to dog identity (dog_fraction). Host-associated variance was detectable at phylum level but became stronger and more interpretable at genus and species levels. Several genera showed high dog_fraction values, including *Weissella*, *Thauera*, *Saccharibacillus*, *Dialister*, *Pediococcus*, *Lactococcus*, and *Campylobacter*, with dog identity explaining up to about 35–60% of abundance variance for selected genera (**Figure 2B**; **Supplementary Table S7**).

Species-level profiles revealed the clearest individualized signatures. Multiple species showed substantial host-associated variance fractions, often reaching about 40–60%, and displayed distinct temporal patterns across dietary stages (**Figure 2B–C**; **Supplementary Table S8A–B**). Some taxa, such as *Lactococcus garvieae*, showed stage-restricted peaks confined to a small subset of dogs, whereas others, including [*Ruminococcus*] *torques* (*Mediterraneibacter*), showed persistent between-dog differences across multiple stages. *Bifidobacterium pseudolongum* showed a third pattern, with a broader stage-associated increase but marked variation in magnitude among individual dogs.

Some dogs consistently showed elevated abundance across multiple host-associated taxa, whereas others showed enrichment for only one focal species or none. These patterns indicate overlapping individualized microbial signatures rather than a single uniform host-associated profile.

Finally, we tested whether maternal background contributed to early microbiome structure by restricting the analysis to Ag1 samples, when puppies were exclusively breastfed. Litter identity explained 10.8% of community variation in ONT 16S profiles and 13.1% in mWGS profiles, with the latter approaching statistical significance (R² = 0.1305, p = 0.062). Thus, maternal/litter background contributed modestly to neonatal community structure, but the stronger longitudinal pattern was persistent host-specific organization across development (**Supplementary Results 2**).

### 3. Virome remodeling accompanies early-life microbiome maturation

To characterize virome dynamics during microbiome maturation, we analyzed viral contigs recovered from Illumina metagenomic co-assemblies. The assembly yielded 1,092,076 contigs, of which 5,109 were classified as viral. After stringent quality filtering, 2,631 high-confidence viral contigs remained, including 2,447 prokaryotic viruses. High-confidence host predictions were obtained for 957 viral contigs corresponding to 191 bacterial genera (**Figure 3A–D**).

**Figure 3.**
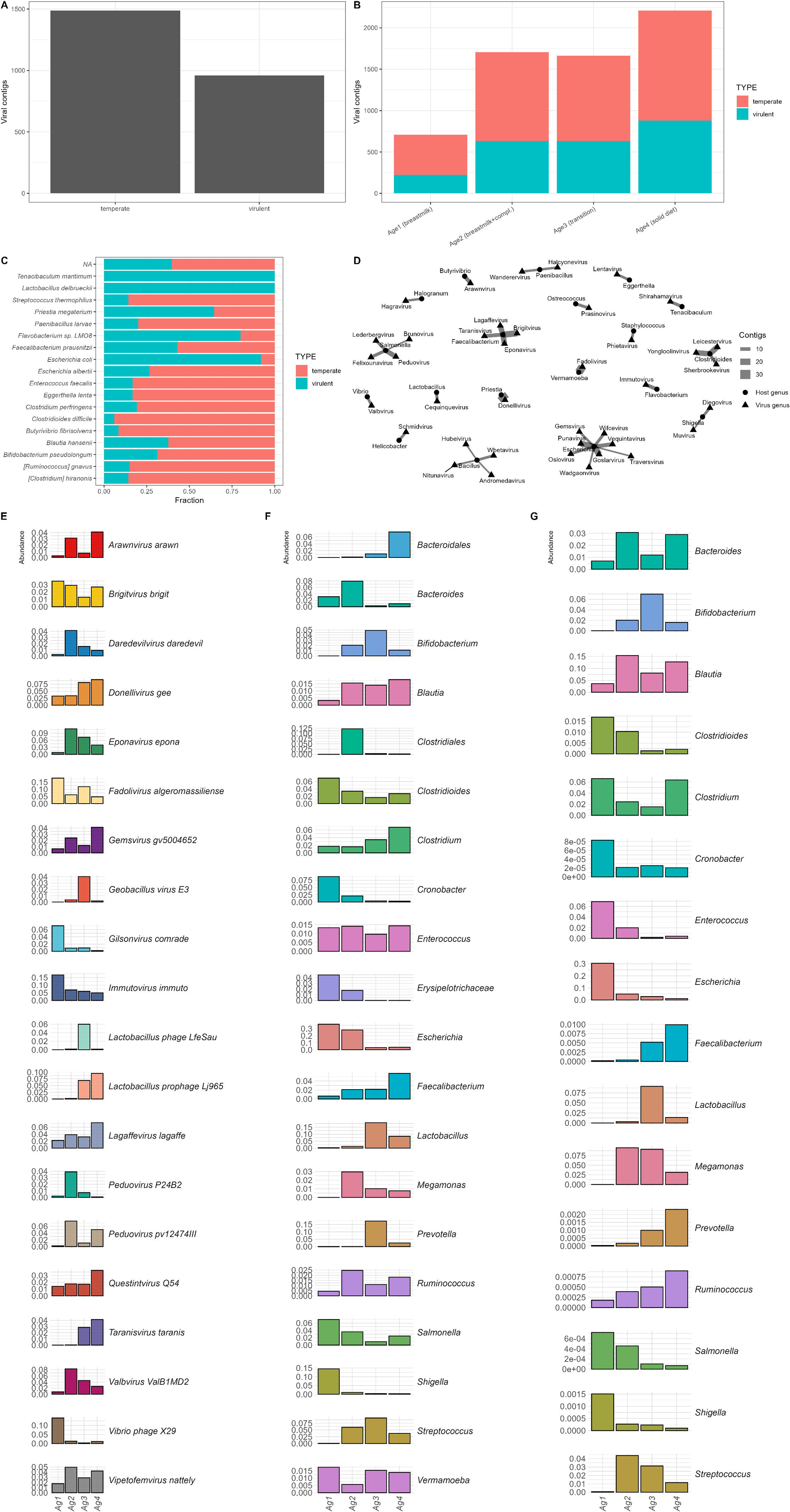
Virome composition, lifestyle distribution, and predicted host associations across early-life dietary stages. **A**, Distribution of predicted viral lifestyles among high-confidence viral contigs. **B**, Relative proportions of temperate and virulent phages across developmental stages. **C**, Distribution of viral lifestyles across predicted bacterial host genera. **D**, Virus–host interaction network based on high-confidence CHERRY predictions. Nodes represent viral and bacterial genera, and edge widths indicate the number of contigs supporting each association. Only interactions supported by at least two contigs and the most connected taxa are shown. **E**, Relative abundance patterns of the 20 most frequent viral species across developmental stages. **F**, Relative abundance of predicted bacterial host genera inferred from CHERRY host predictions. **G**, Relative abundance of bacterial genera based on whole-metagenome Kraken profiling.

Virome richness increased progressively across dietary development. Using a threshold of >10 mapped reads per group, we detected 807, 1,955, 1,850, and 2,434 unique viral contigs in Ag1–Ag4, respectively, with the highest richness observed under established solid diet conditions (**Figure 3B**). Consistent with this pattern, the proportion of reads mapping to viral contigs increased from 16.5% to 30.2% across developmental stages.

Temperate phages dominated the virome across all age groups, indicating that lysogenic interactions represent a major component of the early-life canine gut virome (**Figure 3A–C**). However, predicted virus–host associations showed substantial heterogeneity across bacterial genera. *Bacteroides* and *Faecalibacterium* were associated predominantly with temperate phages, whereas *Enterococcus* and *Escherichia* showed more balanced associations with both temperate and virulent viruses (**Figure 3C**).

Viral community composition also changed across dietary stages. Several viral taxa, including *Araarivirus arawn*, *Gemsvirus gv5004652*, and *Quenstivirus Q54*, increased progressively with age and reached highest abundance in Ag4, whereas others, including *Immutovirus immuto* and *Vibrio* phage X29, were enriched during early breastfeeding stages and declined later (**Figure 3E**).

Predicted bacterial host profiles derived from viral host assignment broadly mirrored the taxonomic maturation patterns observed in whole-community metagenomes. Host-associated genera linked to later-stage microbiome maturation, including *Bacteroides*, *Faecalibacterium*, *Ruminococcus*, and *Prevotella*, increased across development, whereas early colonizers such as *Escherichia* were enriched during breastfeeding stages (**Figure 3F–G**). Bacterial and viral community changes therefore appeared coupled during early-life dietary maturation (**Supplementary Results 3**).

### 4. Functional maturation accompanies taxonomic succession

We next asked whether the taxonomic succession observed across dietary stages was accompanied by functional restructuring of the gut microbiome. KEGG ortholog profiles derived from mWGS data were analyzed with DESeq2, followed by pathway-level over-representation analysis of significantly differentially abundant KOs (FDR < 0.05). Directionality was assessed by summarizing KO-level log2 fold changes within enriched pathways (**Figure 4A–D**; **Supplementary Tables S9–S10**).

**Figure 4.**
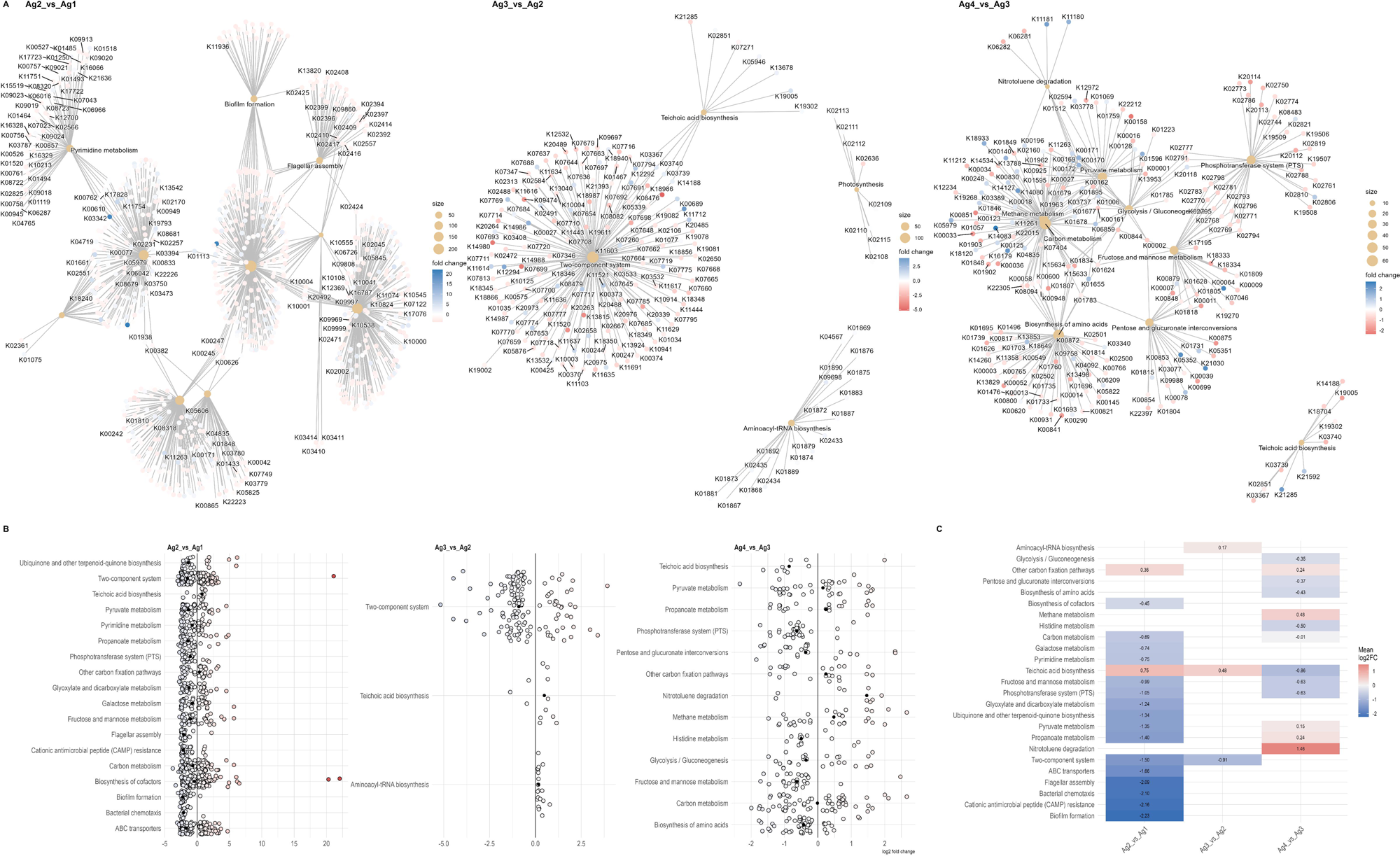
Functional restructuring of the gut microbiome across dietary transitions. **A**, Networks of enriched KEGG pathways and associated KOs across consecutive developmental transitions. Large nodes represent pathways and small nodes represent KOs. Node size reflects KO count per pathway, node color indicates log2 fold change, and edges denote KO–pathway associations. **B**, Directional pathway changes across transitions. Each point represents an enriched pathway, and the vertical line indicates no change. Positive values denote enrichment in the later stage. **C**, Integrated heatmap of mean pathway log2 fold changes across transitions. Red indicates enrichment and blue indicates depletion in the later stage; cell values show mean log2 fold changes.

The strongest functional shift occurred during the Ag1→Ag2 transition, corresponding to the onset of dietary diversification. Enriched pathways included flagellar assembly, biofilm formation, bacterial chemotaxis, and two-component signal transduction, alongside central metabolic pathways such as carbon metabolism, biosynthesis of cofactors, and quinone biosynthesis (**Figure 4A**). Several of these pathways showed strong enrichment, including flagellar assembly and biofilm formation (approximately 2.0-fold enrichment), bacterial chemotaxis (about 1.9-fold), and two-component systems (about 1.36-fold).

Subsequent transitions showed progressively reduced pathway-level turnover. During Ag2→Ag3, enriched functions were concentrated mainly in regulatory and cell-envelope–associated processes, including two-component systems and teichoic acid biosynthesis. By Ag3→Ag4, enriched pathways were fewer and more focused on substrate uptake and central metabolism, including the phosphotransferase system (PTS), which showed approximately 3.0-fold enrichment (**Figure 4A–C**).

Directionality analysis further supported a staged maturation pattern. Colonization-associated pathways in the Ag1→Ag2 comparison, including flagellar assembly and biofilm formation, were biased toward higher abundance in Ag1, whereas later-stage enriched pathways showed smaller and more pathway-specific shifts (**Figure 4B–C**). Early dietary diversification was therefore associated with broad functional reorganization, followed by regulatory refinement and targeted metabolic adjustment under solid diet conditions (**Figure 4D**; **Supplementary Results 4**).

### 5. Matched ONT–mWGS profiling reveals platform-specific taxonomic biases

We next benchmarked ONT full-length 16S taxonomic profiles against matched Illumina mWGS profiles to quantify platform effects and evaluate whether 16S rRNA gene copy-number normalization improved agreement with metagenomic profiles. Community-level PCoA based on Bray–Curtis dissimilarity showed that samples clustered primarily by sequencing strategy rather than dietary stage, with mWGS profiles clearly separated from both raw and rrnDB-normalized ONT 16S profiles (**Figure 5A**).

**Figure 5.**
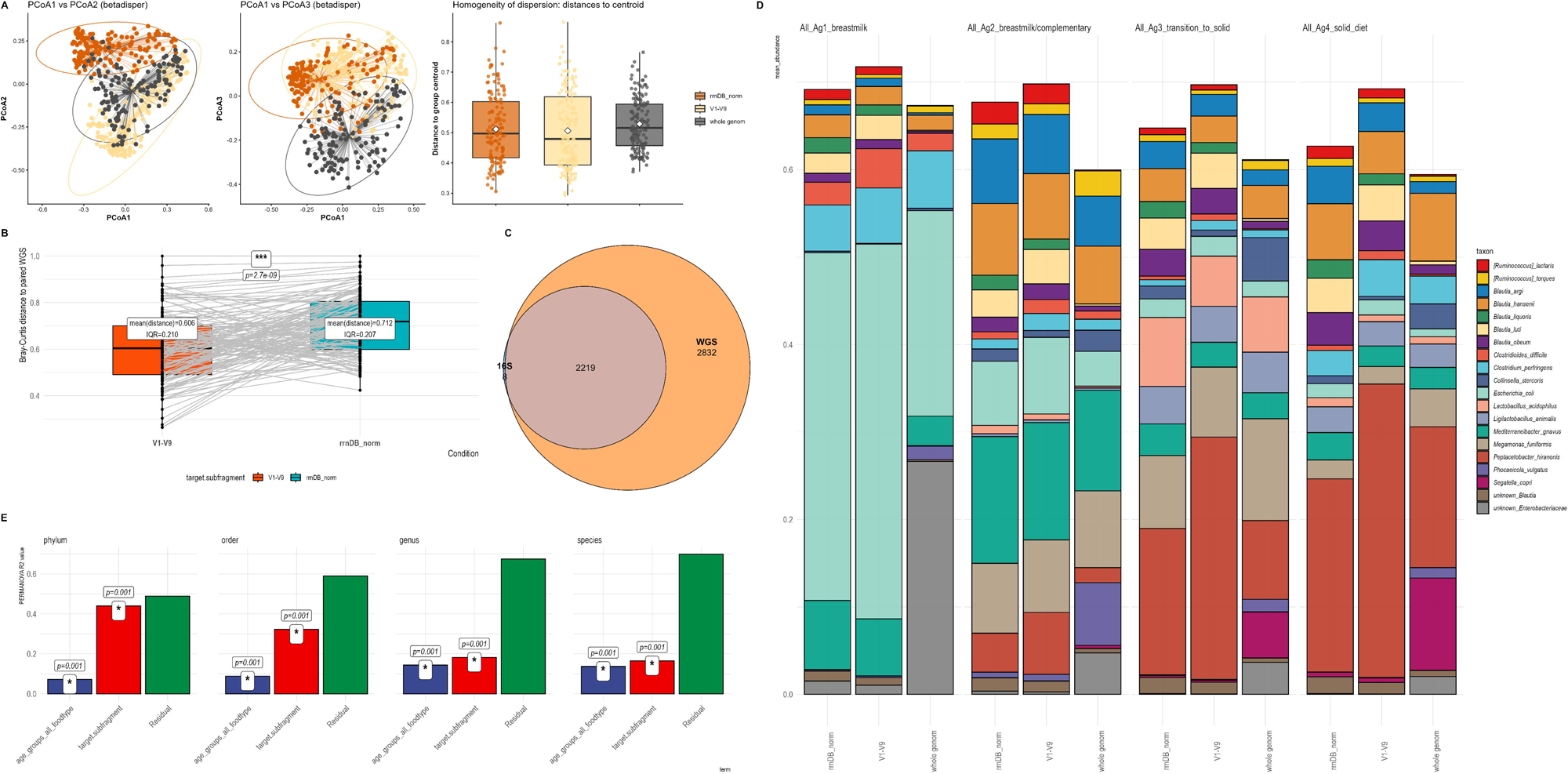
Platform-dependent taxonomic profiles and effects of 16S copy-number normalization. **A**, PCoA of Bray–Curtis dissimilarities comparing mWGS, ONT V1–V9, and rrnDB-normalized V1–V9 taxonomic profiles at species resolution. Left and middle panels show PCoA1–2 and PCoA1–3 projections; ellipses denote group dispersion and line segments connect samples to group centroids. The right panel shows distances to centroids, quantifying dispersion across sequencing strategies. **B**, Paired Bray–Curtis distances between ONT V1–V9 and matched mWGS profiles before and after rrnDB-based copy-number normalization. Points represent samples, lines connect paired measurements, and boxplots summarize distributions. ***P < 0.001, paired test. **C**, Euler diagram showing overlap of species detected by ONT V1–V9 and mWGS. **D**, Mean relative abundances of the top 20 taxa across developmental stages and profiling approaches. Bars represent developmental stages; colors indicate taxonomic identity. **E**, PERMANOVA-derived variance partitioning across taxonomic levels. Bars show R² values for age group and sequencing method; residual variance indicates unexplained heterogeneity. Asterisks denote significant effects.

Stage-associated compositional gradients were still visible across sequencing strategies. Early stages were dominated by Pseudomonadota, whereas later stages showed increased Bacillota, Bacteroidota, and Actinomycetota, although the magnitude of these shifts differed between mWGS, raw ONT 16S, and rrnDB-normalized profiles (**Figure 5B**).

At community level, rrnDB normalization did not improve agreement with matched mWGS profiles. Non-normalized ONT V1–V9 profiles showed a mean Bray–Curtis distance of 0.606 to matched mWGS profiles, whereas rrnDB-normalized profiles showed a higher mean distance of 0.712 (paired Wilcoxon p = 2.7 × 10CC; **Figure 5C**). This pattern remained when analyses were restricted to taxa with direct rrnDB copy-number entries.

Across all samples, mWGS recovered a broader bacterial species repertoire than ONT 16S. A total of 5,059 species were detected, including 2,832 detected only by mWGS, 8 detected only by 16S, and 2,219 shared between methods (**Figure 5D**).

PERMANOVA confirmed that both sequencing method and dietary stage significantly contributed to compositional variation. Method effects were strongest at higher taxonomic ranks, decreasing from R² = 0.44 at phylum level to R² = 0.165 at species level, whereas age-associated variance increased from R² = 0.072 to R² = 0.137 across the same ranks (**Figure 5E**; **Supplementary Table S11**).

Long-read 16S profiles therefore retained developmental signal, but cross-platform comparisons were dominated by methodological differences. rrnDB normalization partially reduced species-level bias for selected taxa, but did not improve global community-level agreement with matched mWGS profiles (**Supplementary Results 5**).

### 6. Long-read 16S captures inference-level functional shifts despite taxonomic bias

Finally, we evaluated whether ONT full-length 16S profiles could recover biologically meaningful functional restructuring despite the strong taxonomic platform effects observed in cross-platform comparisons. PICRUSt2-based functional predictions derived from ONT profiles were benchmarked against matched mWGS-derived KEGG ortholog profiles across 142 paired samples using three preprocessing workflows: default NanoCLUST, curated NanoCLUST after strand correction and chimera removal, and NanoASV with stringent placement thresholds.

Read retention differed substantially among workflows (**Figure 6A**; **Supplementary Table S12**). Default NanoCLUST retained 0.47–0.57 of reads across developmental stages, whereas curated NanoCLUST increased retention to 0.57–0.72. NanoASV achieved the highest retention, ranging from 0.78 to 0.91 across age groups.

**Figure 6.**
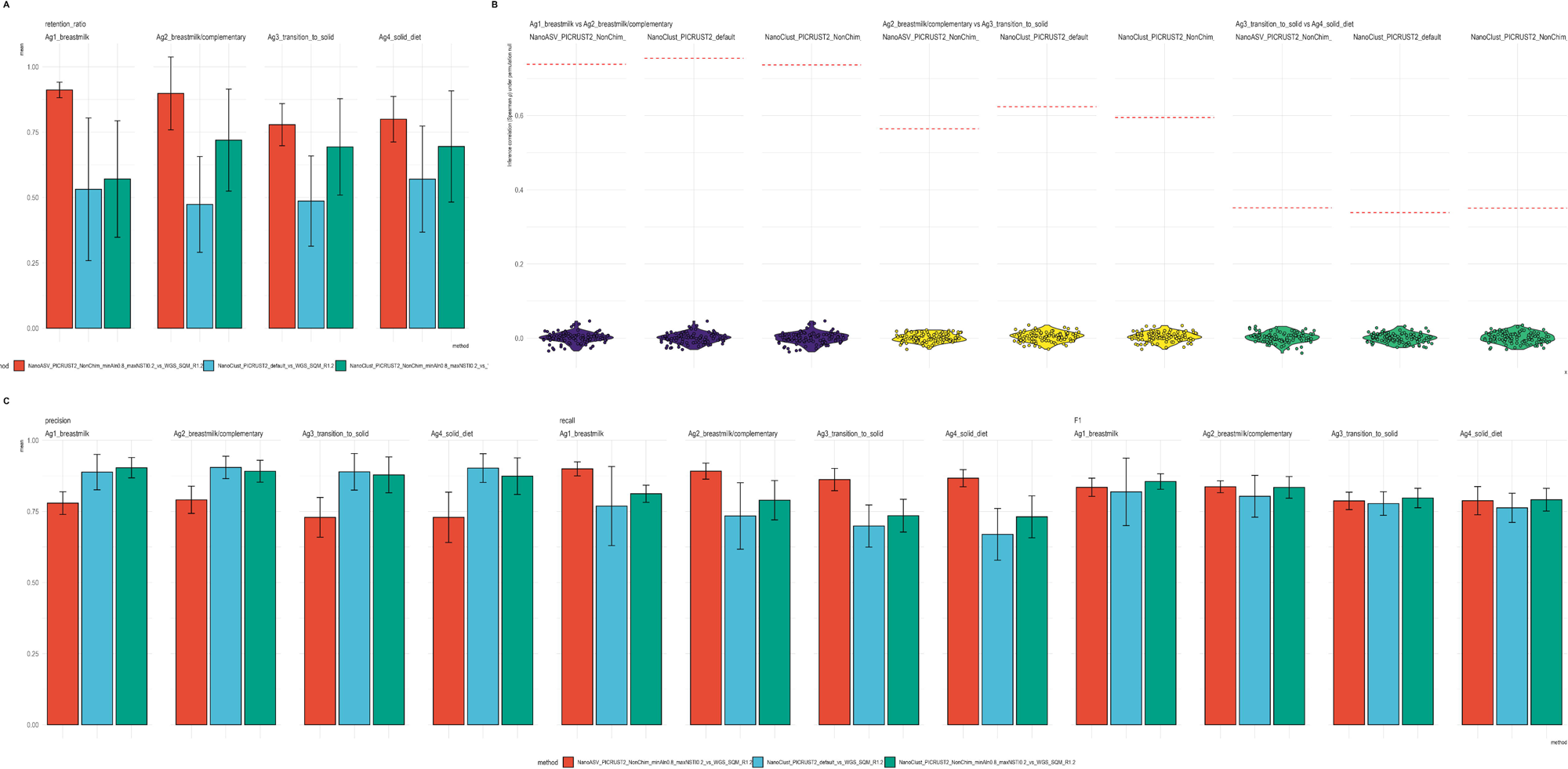
Functional inference from long-read 16S profiles and concordance with metagenomic functional restructuring. **A**, Proportion of ONT reads retained after PICRUSt2 placement and NSTI filtering across preprocessing workflows and developmental stages. Bars represent mean ± SD across samples. **B**, Inference-level concordance between PICRUSt2-derived and mWGS-derived functional contrasts across developmental transitions, evaluated using signed log10-transformed KO-level statistical effect vectors from Wilcoxon rank-sum tests. Points represent permutation-derived null distributions, and dashed red lines indicate observed correlations. **C**, KO detection performance across preprocessing workflows. Precision, recall, and F1 scores are shown for default NanoCLUST, curated NanoCLUST, and NanoASV workflows across developmental stages.

We next evaluated inference-level concordance between PICRUSt2-derived and mWGS-derived functional changes using the contrast-based framework of Sun et al. (2020). Signed log10-transformed KO-level statistical effect vectors from developmental contrasts (Ag1→Ag2, Ag2→Ag3, Ag3→Ag4) were correlated between inferred and metagenomic profiles and compared against permutation-derived null expectations (**Figure 6B**; **Supplementary Table S14**).

Across all workflows and developmental contrasts, observed correlations substantially exceeded null expectations. Spearman correlations ranged from 0.338 to 0.755 and Pearson correlations from 0.408 to 0.776, whereas permutation-based null correlations remained centered near zero (|ρ_perm| < 0.01; one-sided empirical p = 0.0099). Concordance was strongest for the Ag1→Ag2 transition and weaker but consistently above null expectations for Ag3→Ag4.

KO detection metrics revealed distinct precision–recall trade-offs among preprocessing strategies (**Figure 6C**; **Supplementary Table S13**). Default NanoCLUST maintained high precision (0.889–0.905) but lower recall (0.670–0.769), whereas NanoASV achieved consistently higher recall (0.862–0.900) at the expense of reduced precision (0.729–0.791). Despite these differing error profiles, F1 scores remained broadly comparable across workflows.

Thus, long-read 16S workflows recovered pathway-level developmental contrasts despite strong platform-dependent taxonomic bias, with preprocessing primarily influencing precision–recall balance rather than overall inference-level concordance (**Supplementary Results 6**).

## Discussion

In this longitudinal study of Pumi puppies, early-life gut microbiome maturation followed a shared age–diet-associated trajectory, but this trajectory unfolded within persistent host-specific microbial structure. Across bacterial, viral, and functional profiles, the strongest developmental changes occurred during the transition from milk feeding toward complementary feeding and solid diet. However, these cohort-level shifts occurred on individualized abundance baselines that persisted across development.

Taxonomically, the strongest compositional restructuring occurred during the transition from breastfeeding toward complementary feeding and solid diet. Early communities were characterized by low diversity and high relative abundance of facultative taxa, particularly *Escherichia coli*, which declined from approximately 23.5% mean relative abundance in Ag1 to 0.9% in Ag4. In parallel, later stages showed expansion of anaerobe-associated taxa including *Segatella copri*, *Peptacetobacter hiranonis*, and *Megamonas funiformis*. These coordinated taxonomic shifts indicate progressive ecological succession from early facultative assemblages toward a more diverse anaerobe-rich community adapted to increasingly complex dietary substrates.

Functional restructuring closely paralleled taxonomic succession. The Ag1→Ag2 transition exhibited the strongest pathway-level signal, including enrichment of motility-, chemotaxis-, biofilm-, and signal-transduction–associated pathways alongside central metabolic functions. Later transitions involved progressively fewer enriched pathways and smaller KO-level shifts, indicating reduced functional turnover as the microbiome approached a more stabilized configuration under solid diet conditions. This supports a staged maturation model, in which early ecological restructuring is followed by regulatory and metabolic refinement.

Beyond developmental stage, host identity emerged as a major determinant of microbiome structure. Across taxonomic levels, PERMANOVA consistently identified dog identity as explaining a larger fraction of compositional variance than age/food group. This pattern was reinforced by taxon-wise CLR-based mixed-effects models that attributed host-associated variance fractions (dog_fraction) reaching about 40–60% for several species.

Importantly, the mixed-effects analysis should be interpreted as variance partitioning rather than classical differential-abundance testing. Rather than identifying taxa differing between predefined host groups, the model estimates how strongly taxon-level abundance variation remains structured by host identity after accounting for developmental stage. These results indicate that microbiome maturation occurs within individualized abundance baselines rather than replacing them.

At coarse taxonomic ranks, host effects were detectable but more diffuse, reflecting the aggregation of multiple lower-level lineages with potentially different temporal behaviors. At genus and especially species resolution, host-associated variance became easier to interpret, revealing taxa whose abundances were consistently individualized across time. Several taxa that changed strongly across age groups nevertheless retained marked between-dog differences, whereas others showed transient peaks restricted to a few hosts. Together, these patterns indicate that developmental change and host individuality were not competing explanations, but parallel components of early-life microbiome maturation.

Maternal background contributed more modestly to early community organization. Restricting analyses to Ag1 samples, litter identity explained 10.8% of variation in ONT profiles and 13.1% in mWGS profiles, with the latter approaching statistical significance. Although these results suggest a limited litter-associated effect during the neonatal stage, the dominant longitudinal pattern remained persistent host-specific organization across development.

The staged transition observed in this cohort parallels ecological principles described in human infant microbiome studies, where facultative anaerobes dominate early communities before progressive expansion of obligate anaerobes following dietary diversification and weaning. At the same time, the mature canine microbiome retains characteristic lineage compositions distinct from humans, including prominent Fusobacteriota and canine-associated Clostridia/Lachnospiraceae assemblages. Thus, the canine system captures conserved principles of early-life microbiome maturation while preserving host-specific ecological structure.

Comparison with previously published canine cohorts further supports the reproducibility of the developmental trajectory observed here. Relative to published neonatal datasets, our Ag1–early Ag2 communities showed a comparatively delayed expansion of *Bacteroidota* and Fusobacteriota, remaining dominated by *Bacillota* and *Pseudomonadota* during early stages. Such differences likely reflect variation in litter structure, maternal microbiota, husbandry conditions, and cohort composition, together with methodological differences between studies. Nevertheless, by Ag4 the microbiome converged toward the *Bacillota*–*Bacteroidota*–*Fusobacteriota*-dominated structure commonly reported in post-weaning and adult canine microbiomes.

A methodological strength of this study is the paired sequencing design, which allowed us to distinguish biological developmental signal from platform-dependent bias. Although developmental trajectories were detectable across methods, sequencing platform explained a substantial proportion of compositional variance and exceeded biological stage effects at higher taxonomic ranks. mWGS recovered a substantially broader species repertoire, whereas most taxa detected by 16S belonged to a shared core also identified in metagenomes. rrnDB-based copy-number normalization produced species-dependent effects: agreement with mWGS improved for some abundant taxa but worsened for others, and community-level Bray–Curtis similarity to mWGS decreased after normalization. Thus, rRNA operon copy-number correction should not be treated as a default improvement for full-length 16S profiles. In this dataset, its effect was taxon-dependent and it did not improve global agreement with matched metagenomes.

Despite strong platform-dependent taxonomic biases, functional inference from long-read 16S profiles showed robust concordance with metagenome-derived pathway shifts. Across all preprocessing workflows, PICRUSt2-derived functional contrasts correlated strongly with mWGS-derived contrasts and consistently exceeded permutation-based null expectations. NanoASV maximized read retention and recall, whereas NanoCLUST-based workflows retained higher precision, but overall inference-level concordance remained broadly stable across preprocessing strategies. These results suggest that long-read 16S sequencing, when coupled with inference-level validation, can recover biologically meaningful pathway-level developmental patterns despite imperfect taxonomic agreement with shotgun metagenomics.

Overall, these data support a model of host-constrained ecological succession in the early-life canine gut (**Figure 7**). Age–diet progression defined the shared direction of maturation, whereas host identity shaped taxon-specific abundance trajectories across that transition. The study also illustrates how shotgun metagenomics, virome profiling, and long-read 16S sequencing can be combined to study microbiome development in longitudinal animal cohorts.

**Figure 7.**
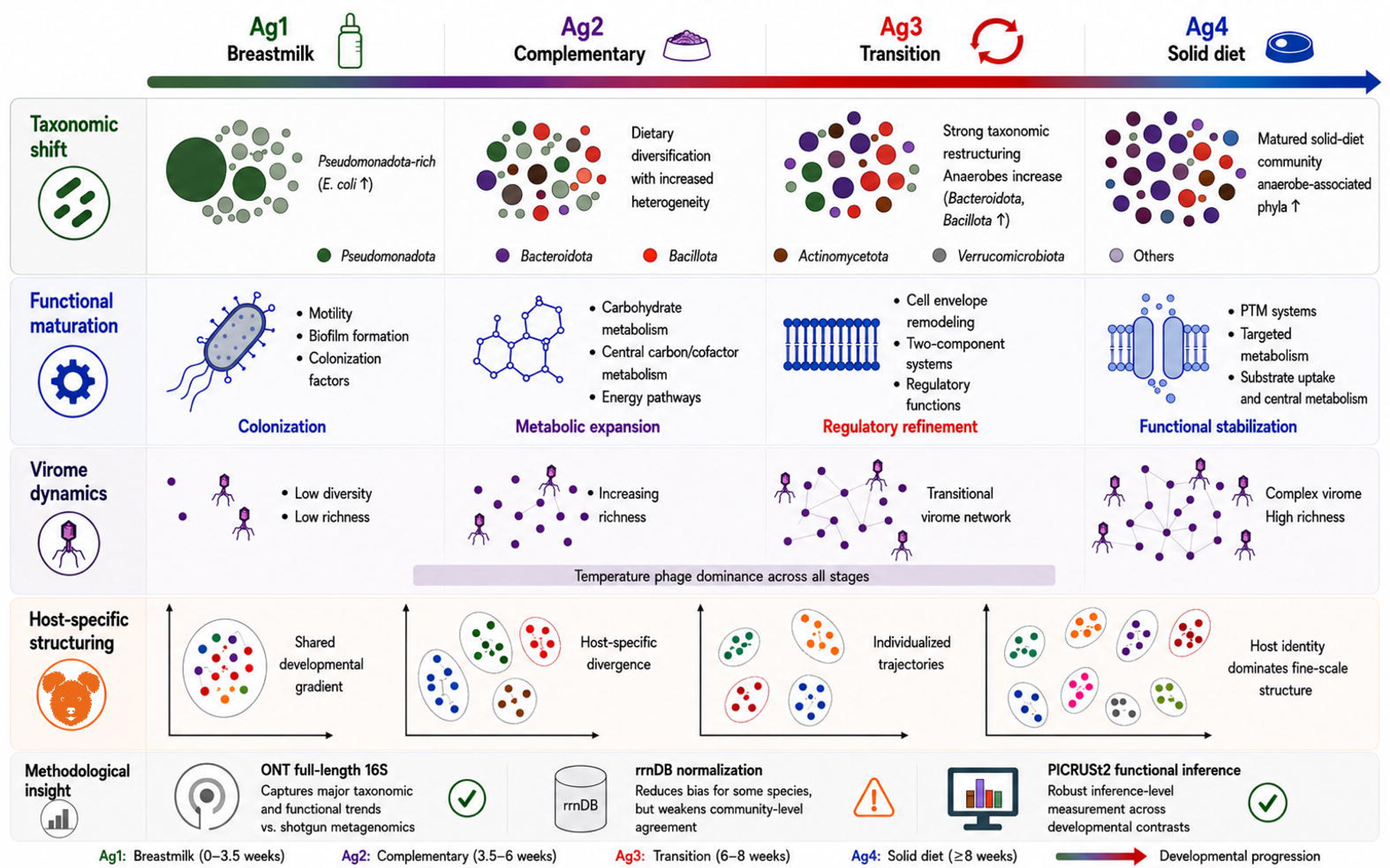
Integrative model of diet-driven microbiome maturation with increasing host-specific structuring and virome complexity in early-life dogs.

## Limitations

Several limitations should be acknowledged. The cohort was modest in size compared with large population-based microbiome studies, although the controlled single-kennel, single-breed design improved internal consistency and reduced environmental heterogeneity. Age and diet could not be fully separated, because each developmental stage corresponded to a defined feeding regime. Virome analyses were based on viral contigs rather than complete viral genomes, and functional predictions from long-read 16S profiles were benchmarked against metagenomes but not validated using metatranscriptomics or metabolomics. Fungal communities were not characterized because of extraction constraints.

## Conclusions

Early-life canine gut microbiome maturation involved coordinated bacterial succession, virome remodeling, and functional restructuring across dietary development. This process followed a shared diet-associated trajectory, but host identity strongly individualized microbial abundance patterns across stages. Cross-platform benchmarking further showed that ONT full-length 16S sequencing retains developmental signal despite substantial taxonomic bias, and that 16S-based functional inference can recover pathway-level contrasts concordant with matched metagenomes. These findings provide biological insight into early-life canine gut ecosystem development and practical guidance for integrating long-read 16S and shotgun metagenomic profiling in longitudinal microbiome studies.

## Methods

### Sample collection and study design

Fecal samples were collected from Pumi dogs representing three litters (B, C, and D) born between 2020 and 2023. In total, 146 fecal samples were obtained across four developmental stages, with each dog contributing a minimum of four samples spanning early-life transitions. Immediately after collection, samples were stored at –20C°C, transported on dry ice within one week, and archived at –80C°C until processing. Comprehensive sample-level metadata, including sample identifiers, dog identity, litter assignment, sex, birth date, sampling date and age–diet group classification, are provided in **Supplementary Table S15**. The overall experimental and bioinformatic workflow is summarized in **Supplementary Figure S2**.

All animals were housed in the same kennel (Baja, Hungary), under uniform environmental and care conditions. Each litter had a distinct sire (none of which resided at the sampling site), while litters B and C shared the same dam. Litter D descended from a female in litter B, creating a vertical maternal lineage across generations. This familial structure allowed structured comparison of age, diet, and maternal background in relation to microbiome development (**Supplementary Figure S3**). The cohort included 11 males and 8 females. Sex was not a significant predictor of microbiome composition in preliminary analyses (PERMANOVA, p > 0.05) and was not included in final models.

Dietary variation among litters served as a natural experimental contrast. The dam of litters B and C was initially fed a commercial dry diet (Brit Care Hypoallergenic Dog Show Champion), but prior to conceiving litter C, she was transitioned to a BARF (Biologically Appropriate Raw Food) diet. She reverted to dry food postpartum due to poor tolerance. The mother of litter D, herself born in litter B, had been maintained on a BARF diet from one year of age onward.

Body weight monitoring. To monitor early postnatal growth as a general health indicator, puppies were weighed weekly during the first four weeks of life. Litter sizes differed (B, n=4; C, n=5; D, n=8), which may influence early growth via nursing competition; weight distributions and individual trajectories are shown in **Supplementary Figure S4**.

### Pedigree structure and genetic relatedness

Litters B and C share a dam but unrelated sires, making them maternal half-siblings (second-degree relatives). Litter D represents the next generation, descended from a female in litter B and a genetically distinct male. As a result, puppies in litter D are nieces and nephews of those in litters B and C. The pedigree thus establishes a quasi-longitudinal framework with descending degrees of relatedness: highest between B and C, intermediate between C and D, and most distant between B and D. One neutered adult dog, distantly related to the maternal lineage of litters B and C, was also included for comparison, adding diversity while maintaining partial genetic continuity (**Supplementary Figure S3**).

### Dietary stage classification

Puppies were grouped into four distinct age–diet stages, based on feeding milestones and breeder-reported dietary records: (1) Ag1 (breastmilk only): ≤3.5 weeks of age, exclusively nursing from the dam. (2) Ag2 (breastmilk + complementary feeding): 3.5–6 weeks, with gradual introduction of solid food alongside milk. (3) Ag3 (transition to solid food): >6 to 8 weeks, characterized by increasing reliance on solid food and declining milk intake. (4) Ag4 (solid diet): ≥8 weeks, complete cessation of nursing due to tooth eruption and exclusive consumption of solid food. This classification framework allowed high-resolution mapping of microbiome transitions across early life.

### DNA extraction

Genomic DNA was extracted from 200 mg of fecal material using the NucleoSpin® DNA Stool Kit (Macherey-Nagel), following the manufacturer’s protocol. DNA was eluted in 100 µL of Buffer SE and stored at –20°C. Sample integrity, purity, and yield were assessed using Qubit fluorometry and Agilent TapeStation.

### Library preparation and sequencing

#### Illumina whole-genome shotgun (mWGS) libraries

For mWGS libraries, 50 ng of purified DNA per sample was processed using the Illumina DNA Prep Kit. The tagmentation and amplification protocols followed the manufacturer’s instructions. Indexed libraries were purified via magnetic bead cleanup and eluted in resuspension buffer (supplied with the kit). Quality control and fragment size distribution were assessed using the Agilent D1000 ScreenTape.

#### ONT 16S rRNA (V1–V9) amplicon libraries

ONT libraries were generated using the 16S Barcoding Kit (SQK-16S024) with 10 ng of high-molecular-weight DNA as input. PCR amplification was performed with LongAmp Taq polymerase under standard cycling conditions (25 cycles). Libraries were purified using AMPure XP beads and resuspended in 10 mM Tris-HCl + 50 mM NaCl for sequencing. Fragment quality was confirmed with the D5000 ScreenTape.

### Bioinformatic Methods

All analyses were conducted in R (v4.3.0 or later).

#### Illumina Shotgun Metagenomic Sequencing and Processing

Illumina paired-end reads were trimmed using Trimmomatic v0.39 (Bolger et al., 2014) to remove adapters and low-quality bases (LEADING:8, TRAILING:8, SLIDINGWINDOW:10:15, MINLEN:30). Host-derived reads were removed by aligning against the CanFam3.1 reference genome using Bowtie2 with the --very-sensitive-local preset (Langmead and Salzberg, 2012). Only unmapped reads were retained for microbiome analyses.

Taxonomic classification of filtered reads was performed using Kraken2 v2.1.3 with the standard database and default parameters (Wood et al., 2019). Both per-read classifications and sample-level summary reports were generated for downstream taxonomic profiling.

For functional profiling, mWGS reads were processed using SqueezeMeta v1.6.5 in co-assembly mode (Tamames and Puente-Sánchez, 2019). After internal quality filtering, reads were co-assembled using MEGAHIT (Li et al., 2015). Open reading frames were predicted using Prodigal in metagenomic mode (Hyatt et al., 2010), and functional annotation was performed with DIAMOND (Buchfink et al., 2015) against KEGG and COG databases. KEGG Orthology (KO) tables were extracted for downstream analyses. Output tables were imported into R using SQMtools (Puente-Sánchez et al., 2020) and integrated into phyloseq objects (McMurdie and Holmes, 2013).

#### ONT Full-Length 16S rRNA Sequencing and Processing

Full-length 16S rRNA (V1–V9) amplicon data generated using ONT were processed with the NanoASV pipeline (Cousson et al., 2025). Clustering and consensus sequence generation were performed within the workflow, and resulting feature tables were exported in BIOM format. To ensure consistent orientation, consensus sequences were aligned against the SILVA SSU database (Chuvochina et al., 2025) using the --orient function of VSEARCH (Rognes et al., 2016). Chimeric sequences were removed using a combined UCHIME-ref (Gold/RDP reference) and UCHIME de novo strategy (Edgar et al., 2011). Taxonomic assignments were derived from the NanoASV output.

#### 16S rRNA Gene Copy Number Normalization

To correct for interspecific variation in ribosomal operon copy number, species-level ONT relative abundances were normalized using rrnDB v5.9 pantaxa statistics (Stoddard et al., 2015). Observed relative abundances were divided by the mean operon copy number reported for each species. For taxa lacking a corresponding rrnDB entry, the global rrnDB mean (about 4.5 copies per genome) was used as a fallback value to minimize systematic bias while retaining taxa with incomplete annotation. Copy-number–normalized profiles were analyzed alongside raw 16S and mWGS-derived taxonomic profiles.

#### Functional Inference from 16S Data

Functional prediction from ONT 16S data was performed using PICRUSt2 (Douglas et al., 2020) under stringent criteria. Only sequences meeting a minimum alignment threshold of 0.8 and a nearest sequenced taxon index (NSTI) cutoff ≤ 0.2 were retained for inference. Predicted KO and KEGG pathway abundances were used for comparative and benchmarking analyses against metagenome-derived annotations. Three ONT preprocessing workflows were compared for functional benchmarking: default NanoCLUST consensus sequences, curated NanoCLUST consensus sequences after strand correction and chimera filtering, and NanoASV-derived ASVs. Downstream PICRUSt2 processing was performed consistently across workflows.

#### Data Integration

All taxonomic and functional abundance matrices—including raw 16S, rrnDB-normalized 16S (Stoddard et al., 2015), mWGS taxonomic profiles, SqueezeMeta-derived KO and pathway tables, and PICRUSt2 predictions—were integrated into unified phyloseq objects (McMurdie and Holmes, 2013). Sample metadata, taxonomic annotations, and feature tables were harmonized to ensure consistent downstream statistical modeling and visualization.

#### Community Dissimilarity and Ordination

Community dissimilarity was quantified using Bray–Curtis distances calculated from normalized relative abundance tables. This metric captures compositional differences while ignoring joint absences and is appropriate for ecological count data.

Ordination analyses were conducted using both non-metric multidimensional scaling (NMDS) and principal coordinates analysis (PCoA) implemented in the vegan R package (Oksanen et al., 2022). NMDS was performed using the metaMDS function with two-dimensional solutions and stress minimization to assess convergence quality. PCoA was conducted via classical multidimensional scaling of Bray–Curtis distance matrices, and the proportion of variance explained by leading axes was calculated from eigenvalues.

#### PERMANOVA: Partitioning Host and Developmental Effects

Permutational multivariate analysis of variance (PERMANOVA) was performed using adonis2 in vegan (v2.6 or later) to quantify the relative contributions of host identity (dog_alias) and age–diet group (age_groups_all_foodtype) to community structure. Analyses were conducted across multiple taxonomic resolutions using normalized relative abundance data.

Sequential (Type I) models were fitted with alternative term orderings (host → age and age → host) to assess order dependence. Marginal (order-independent) effects were estimated using the by = “margin” option. Host–developmental PERMANOVA models were evaluated with 999 permutations without stratification. For platform-comparison PERMANOVA analyses, marginal effects were tested using permutations restricted within host identity to account for repeated matched profiles. For each factor, R², F statistics, and permutation-based P values were reported. Residual variance represented unexplained community heterogeneity

#### Differential Abundance Analysis of Species

Differential abundance analyses were conducted using MaAsLin2 (Mallick et al., 2021) under a linear modeling framework (analysis_method = “LM”) with centered log-ratio (CLR) normalization and no additional transformation. For profiling method comparisons (raw 16S, rrnDB-normalized 16S, mWGS), target.subfragment was modeled as a fixed effect and sample_ID as a random intercept. mWGS was specified as the reference category. Feature filtering thresholds were set at minimum abundance 1×10CC, minimum prevalence 0.2, and minimum variance 0.01.

For stage-associated taxa (mWGS-based analyses), age_groups_all_foodtype was included as a fixed effect and dog_alias as a random intercept. Features were filtered at minimum abundance 1×10CC and minimum prevalence 0.1. Pairwise contrasts between consecutive developmental stages (Ag1→Ag2, Ag2→Ag3, Ag3→Ag4) were obtained by iterative re-referencing of the categorical predictor. Multiple testing correction was performed using the Benjamini–Hochberg false discovery rate (FDR < 0.05).

### Identification, annotation, and downstream analysis of viral contigs

Viral contigs were identified from the Illumina metagenomic co-assembly using the PhaBOX software (Shang et al., 2023). Contigs were screened by the PhaBOX viral identification module, PhaMer (Shang et al., 2022a), which provided contig-level viral scores and confidence labels (PhaMerScore and PhaMerConfidence). Only contigs with a length of at least 3 kb were retained for downstream analyses.

To select high-confidence viral contigs, an additional filtering step was applied. Only contigs predicted as viral with a PhaMerScore of at least 0.90 were retained. For host prediction analyses, the dataset was further restricted to contigs with a CHERRYScore of at least 0.80, excluding entries labeled as “filtered” in the CHERRY output. (Shang and Sun, 2022).

Viral taxonomic classification was performed using the PhaGCN module (Shang et al., 2021), while host prediction was conducted using the CHERRY module. Viral lifestyle classification as temperate, virulent, or unknown was determined using the PhaTYP module (Shang et al., 2022b).

To estimate contig-level abundance, raw metagenomic reads were mapped back to the assembled contigs using Minimap2 (Li, 2018). Contig-level read counts were used to estimate viral relative abundances across samples and developmental stages.

#### Host-Associated Variance Estimation

To quantify host-specific effects independently of developmental stage, we fitted taxon-wise linear mixed-effects models to CLR-transformed abundance data using lme4 in R (Bates et al., 2015). This analysis was conceptually motivated by the variance-partitioning framework of Sweeny et al. (2023), but was implemented here as a CLR-based linear mixed-effects workflow rather than as a direct reproduction of the original GLMM or Bayesian formulations.

For each taxon, abundance counts were transformed using the centered log-ratio (CLR) after addition of a pseudocount of 0.5. Taxa present in fewer than 10% of samples were excluded before model fitting. Each retained taxon was then modeled separately as: CLR abundance ∼ age_groups_all_foodtype + (1 | dog_alias), where age_groups_all_foodtype was treated as a fixed effect and dog_alias as a random intercept. This formulation quantified the extent to which taxon abundance variation was explained by stable between-dog differences after accounting for developmental stage.

Variance components were extracted from each fitted model, and host association was summarized as: dog_fraction = dog_variance / (dog_variance + residual_variance), where dog_variance denotes the random-effect variance attributable to dog identity and residual_variance denotes the remaining unexplained variance. Higher dog_fraction values therefore indicate stronger host-associated structuring of abundance for a given taxon.

Models were fitted across multiple taxonomic resolutions, including phylum, order, genus, and species. Resulting taxa were ranked by dog_fraction, and selected high-effect taxa were visualized using dog-level abundance trajectories across age groups. Because this analysis is based on variance decomposition of CLR-transformed linear mixed-effects models, the resulting dog_fraction values should be interpreted as host-associated variance fractions rather than as direct measures of differential abundance.

#### Functional Pathway Enrichment Analysis

Differential KO abundance between consecutive age groups was assessed using DESeq2 (Wald test) (Love et al., 2014) applied to raw KO count matrices. KOs with Benjamini–Hochberg–adjusted P values (FDR) < 0.05 were used as input for KEGG pathway over-representation analysis (ORA) implemented in clusterProfiler (Yu et al., 2012). Enrichment testing was performed against the universe of all KOs detected in the dataset, and pathways were considered significant at FDR < 0.05.

#### Functional Benchmarking Between 16S and Metagenomics

To evaluate 16S-based functional inference, PICRUSt2-predicted KO profiles from ONT full-length 16S data were benchmarked against matched mWGS-derived KO profiles generated by SqueezeMeta. Analyses were restricted to 142 matched ONT–mWGS sample pairs. Three ONT preprocessing workflows were compared: default NanoCLUST consensus sequences, curated NanoCLUST consensus sequences after strand correction and chimera filtering, and NanoASV-derived ASVs. Downstream PICRUSt2 processing and benchmarking were applied consistently across workflows.

Benchmarking included three complementary analyses. First, read retention was calculated as the fraction of ONT reads represented by ASVs or consensus sequences retained after PICRUSt2 placement and filtering. Second, KO detection accuracy was evaluated by comparing predicted and mWGS-observed KO presence, summarized using precision, recall, and F1 scores. Third, inference-level concordance was assessed by performing KO-wise differential testing independently on PICRUSt2 and mWGS profiles across consecutive developmental contrasts and correlating signed log10-transformed p-value vectors between platforms. Observed correlations were compared with permutation-derived null distributions.

#### Statistical Analysis and Visualization

All statistical analyses were performed in R (R Core Team, 2023). PERMANOVA was conducted with 999 permutations. Differential abundance and enrichment analyses were corrected for multiple testing using the Benjamini–Hochberg procedure. Data visualization was performed using ggplot2 (Wickham, 2016), phyloseq (McMurdie and Holmes, 2013), vegan (Oksanen et al., 2022), and ComplexHeatmap (Gu et al., 2016), together with custom R scripts.

### Availability of Supporting Data

All sequencing data generated in this study can be found in the European Nucleotide Archive under the accession number: PRJEB75762

## Supporting information

Supplementary Results

## Author Contributions

**TJ** carried out DNA purification, ONT and Illumina library preparation, analyzed the data and drafted the manuscript. **GG** analyzed the data. **MA** carried out DNA purification and ONT library preparation. **ÁD** participated in nucleic acid isolation and ONT sequencing. **ZC** participated in nucleic acid isolation and Illumina sequencing. **BK** analyzed the data and drafted the manuscript. **ZB** participated in data analysis, collected the samples and drafted the manuscript. **DT** conceived and designed the experiments, analyzed the data, collected the samples, drafted the manuscript and coordinated the project.

## Ethical approval

In accordance with institutional and national regulations, formal animal ethics approval was not required for this work. The study involved only noninvasive fecal sampling during routine husbandry without any changes to housing, diet, or veterinary care; samples were provided voluntarily by the breeder. All procedures complied with national animal welfare regulations.

## Abbreviations

ASV: amplicon sequence variant
BARF: biologically appropriate raw food
FDR: false discovery rate
GLMM: generalized linear mixed model
KO: KEGG Ortholog
KEGG: Kyoto Encyclopedia of Genes and Genomes
LRS: long-read sequencing
mWGS: metagenomics whole genome sequencing
NMDS: non-metric multidimensional scaling
ONT: Oxford Nanopore Technologies
ORA: over-representation analysis
PCoA: principal coordinates analysis
PERMANOVA: permutational multivariate analysis of variance
PICRUSt2: Phylogenetic Investigation of Communities by Reconstruction of Unobserved States 2
SRS: short-read sequencing
V1-V9: hypervariable regions 1 to 9 of the 16S rRNA gene

## Acknowledgements

This project was supported by the Hungarian Academy of Sciences, Momentum Grant, LP2020-8/2020 to DT and by the National Research, Development and Innovation Office grants: FK 142676 to DT, K 142674 and ADV 152705 to ZB. The APC fee was covered by the University of Szeged, Open Access Fund: 8294, to DT. The authors would like to express their gratitude to Ildikó Abonyi, the owner of the Duna-Menti Dumás Kennel (Baja, Hungary; https://www.pumikennel.eu/), for collecting the samples.

## Supplementary Figure legends

**Supplementary Figure S1. Age-dependent bacterial read counts across sequencing platforms.** Total bacterial read counts obtained from ONT full-length 16S (V1–V9) and Illumina mWGS datasets across developmental stages (Ag1–Ag4). Bars represent cumulative read counts per developmental stage.

**Supplementary Figure S2. Experimental and bioinformatic workflow of the study.** Overview of the longitudinal study design and analytical workflow. Fecal samples collected across four dietary stages were processed using paired Illumina shotgun metagenomics and Oxford Nanopore full-length 16S rRNA sequencing (V1–V9). Downstream analyses included taxonomic profiling, virome characterization, host-associated variance analysis, rrnDB normalization benchmarking, and PICRUSt2-based functional inference benchmarking against matched metagenomes.

**Supplementary Figure S3. Puppy body weight development across litters. A**, Body weight distributions of puppies from litters B, C, and D across the first four postnatal weeks. Boxplots show median, interquartile range, and individual variation within each litter and week. **B**, Longitudinal body weight trajectories of individual puppies across the first four postnatal weeks. Each line represents one dog.

**Supplementary Figure S4. Pedigree structure and sampling timeline of the Pumi cohort.** Pedigree relationships among the three sampled litters collected between 2020 and 2023. Litters B and C share a common dam but have unrelated sires, whereas litter D descends from a female in litter B and a genetically distinct sire, establishing a multi-generational maternal lineage. Colored silhouettes denote sampled individuals by litter, black silhouettes indicate adult dogs, and grey silhouettes indicate unsampled individuals. Sex is shown below each dog symbol. The right panel summarizes the sampling interval across cohorts.

## Legends to Supplementary Tables

**Supplementary Table S1. Per-sample sequencing summary across platforms.** Per-sample sequencing summary for ONT full-length 16S and Illumina mWGS datasets, including total read counts, classified and unclassified reads, bacterial and viral read counts, corresponding read fractions, and associated sample and host metadata.

**Supplementary Table S2. Comprehensive taxonomic read count matrix across sequencing strategies.** Unified taxonomic read count table comprising Illumina mWGS, raw ONT full-length 16S (V1–V9), and rrnDB-normalized ONT 16S profiles across all samples. Rows represent hierarchical bacterial taxa from phylum to species level, and columns correspond to individual samples and sequencing strategies.

**Supplementary Table S3. Pairwise Bray–Curtis dissimilarity matrix across samples.** Pairwise Bray–Curtis dissimilarities calculated from mWGS-derived species-level relative abundance profiles across all available mWGS samples.

**Supplementary Table S4. Full MaAsLin2 results for developmental contrasts.** Complete MaAsLin2 output across taxonomic ranks for consecutive developmental contrasts. The table includes coefficient estimates, standard errors, raw p-values, false discovery rate–adjusted q-values, taxonomic rank, and contrast specification for all tested taxa.

**Supplementary Table S5A. Mean relative abundances of dominant bacterial phyla across developmental stages.** Mean mWGS-derived relative abundances of the 20 most abundant bacterial phyla across Ag1–Ag4.

**Supplementary Table S5B. Mean relative abundances of dominant bacterial orders across developmental stages.** Mean mWGS-derived relative abundances of the 20 most abundant bacterial orders across Ag1–Ag4.

**Supplementary Table S5C. Mean relative abundances of dominant bacterial species across developmental stages.** Mean mWGS-derived relative abundances of the 20 most abundant bacterial species across Ag1–Ag4.

**Supplementary Table S6. PERMANOVA models across taxonomic levels.** Bray–Curtis–based PERMANOVA models evaluating the effects of dog identity and age/food group on microbiome composition across phylum-, order-, genus-, and species-level profiles. Full and marginal model results are shown, including R² values, F-statistics, permutation-based p-values, and model terms.

**Supplementary Table S7. Host-associated variance components from generalized linear mixed models.** Taxon-level generalized linear mixed model results across taxonomic ranks. Relative abundance was modeled with dog identity included as a random effect. Host association is summarized as the fraction of variance explained by dog identity together with model estimates, standard errors, and significance statistics.

**Supplementary Table S8A. Dog-resolved abundance trajectories of representative host-associated species.** mWGS-derived mean relative abundances per dog across developmental stages (Ag1–Ag4) for representative host-associated species shown in Figure 2C.

**Supplementary Table S8B. Dog-resolved abundance trajectories of representative host-associated phyla.** mWGS-derived mean relative abundances per dog across developmental stages (Ag1–Ag4) at phylum level.

**Supplementary Table S9. KEGG ortholog count matrix.** Raw KEGG ortholog (KO) count matrix derived from mWGS-based SqueezeMeta annotations. Rows correspond to KOs and columns correspond to samples. These counts were used as input for DESeq2 differential abundance analysis.

**Supplementary Table S10. KEGG pathway over-representation analysis results.** Over-representation analysis results for KEGG pathways across consecutive developmental contrasts (Ag2 vs Ag1, Ag3 vs Ag2, Ag4 vs Ag3). The table includes pathway identifiers, gene ratios, background ratios, fold enrichment, adjusted p-values, and associated KOs contributing to each enriched pathway.

**Supplementary Table S11. Species-level PERMANOVA for platform comparison.** Species-level Bray–Curtis–based PERMANOVA models partitioning the effects of sequencing method and developmental stage on taxonomic composition. Marginal models were computed with permutations restricted within host identity.

**Supplementary Table S12. Read retention efficiency across preprocessing workflows.** Mean ± SD PICRUSt2 read retention ratios across preprocessing workflows (default NanoCLUST, curated NanoCLUST, NanoASV) and developmental stages.

**Supplementary Table S13. KO detection metrics across preprocessing workflows and developmental stages.** Mean precision, recall, and F1 scores (± SD) comparing PICRUSt2-predicted KEGG ortholog profiles with matched mWGS-derived annotations across preprocessing workflows and developmental stages (Ag1–Ag4).

**Supplementary Table S14. Inference-level concordance between PICRUSt2 and mWGS functional profiles.** Median Spearman correlation coefficients between signed log10-transformed KO-level differential testing vectors derived from PICRUSt2 and mWGS profiles across biological contrasts and preprocessing workflows. Empirical significance was assessed using permutation-derived null distributions.

**Supplementary Table S15. Sample metadata.** Comprehensive metadata table for all fecal samples, including sample identifiers, dog identity, litter assignment, sex, birth date, sampling date, and associated developmental stage information used in downstream analyses.

