## Supplementary Results for "Early-life canine gut microbiome maturation follows a shared age–diet trajectory within persistent host-specific structure"

**Supplementary Results 1. Detailed taxonomic restructuring across dietary stages**

Quantitative Bray–Curtis summaries supported the ordination and clustering patterns shown in **Figure 1A–B**. Early breastmilk-stage samples were most distinct from later solid-diet samples, while intermediate feeding stages showed transitional community configurations. The largest between-group dissimilarity was observed between Ag1 and Ag4, consistent with progressive age–diet-associated restructuring rather than abrupt community replacement.

MaAsLin2 analysis identified widespread age-associated differences across phylum, order, genus, and species levels. The Ag2→Ag3 transition showed the highest number of significantly associated taxa across all taxonomic ranks, whereas Ag1→Ag2 and Ag3→Ag4 showed fewer significant changes. Directionality differed across transitions, with Ag1→Ag2 dominated by taxa enriched in Ag1, Ag2→Ag3 showing a more balanced pattern of depletion and enrichment, and Ag3→Ag4 showing a modest predominance of taxa enriched in Ag4.

At higher taxonomic ranks, age-associated differences involved dominant lineages rather than rare taxa. Approximately 70–75% of the 20 most abundant phyla and 75–85% of the 20 most abundant orders were significantly associated with at least one developmental transition. These shifts indicate that the observed beta-diversity structure was driven by coordinated changes among major bacterial groups.

At species level, 15 of the 20 most abundant species were significantly age-associated. E. coli showed strong negative associations during early transitions, with MaAsLin coefficients of −2.36 for Ag1→Ag2 and −3.07 for Ag2→Ag3, consistent with its decline from 23.5% in Ag1 to 0.9% in Ag4. In contrast, S. copri increased from 0.35% in Ag2 to 10.5% in Ag4 and showed a strong positive association in Ag2→Ag3 (coef = +5.83, q = 1.28 × 10⁻¹⁸). P. hiranonis increased from 1.7% in Ag2 to 16.0% in Ag4 and was also strongly associated with Ag2→Ag3 (coef = +4.89, q = 9.2 × 10⁻¹³). Additional age-associated species included Megamonas funiformis, which increased from 0.036% in Ag1 to 8.76% in Ag2, and Bifidobacterium pseudolongum, which increased from 0.0005% in Ag1 to 4.8% in Ag3. For these exemplar taxa, the direction and magnitude of MaAsLin2 coefficients were concordant with observed relative abundance changes across dietary stages.

### Supplementary Results 2. Host-associated structuring of early-life microbiome trajectories

PERMANOVA analyses were used to compare the relative contribution of dog identity and age/food group to microbiome composition across taxonomic levels. Across phylum-, genus-, and species-level profiles, dog identity consistently explained a larger fraction of variation than age/food group. At phylum level, total explained variance was lower and overlap between host and age effects was more substantial, whereas at genus and species levels the host-associated component became more pronounced. These results indicate that host identity is a major organizing axis of microbiome composition, particularly at finer taxonomic resolution.

Taxon-wise CLR-based mixed-effects models further supported this pattern. For each taxon, CLR-transformed abundance was modeled as a function of age group, with dog identity included as a random intercept. Host association was summarized as the fraction of modeled variance attributable to dog identity (dog_fraction). At phylum level, host-associated variance was detectable but comparatively moderate, involving phyla such as *Fibrobacterota*, *Acidobacteriota*, *Actinomycetota*, *Balneolota*, and *Campylobacterota*. Even among dominant phyla such as *Bacillota* and *Bacteroidota*, changes across dietary stages were not uniform across the cohort, with individual dogs showing heterogeneous or opposing trajectories.

At genus level, dog identity explained a large fraction of abundance variance for selected taxa, reaching approximately 35–60% for genera including Weissella, Thauera, Saccharibacillus, Dialister, Pediococcus, Lactococcus, and Campylobacter. These effects were typically driven by asynchronous, dog-specific dynamics rather than uniform cohort-wide changes.

Species-level analyses provided the clearest host-associated signatures. Several species showed dog-associated variance fractions (dog_fraction) of about 40–60%. *Lactococcus garvieae* displayed stage-restricted, individual-specific enrichment, mainly during the breastmilk/complementary feeding period, with high abundance confined to a small subset of dogs. Similar temporally localized patterns were observed for *Campylobacter* *upsaliensis* and *Bacteroides fragilis*. In contrast, [*Ruminococcus*] *torques* (*Mediterraneibacter*) showed persistent between-dog differences across multiple stages, while *Bifidobacterium* *pseudolongum* increased during the transition toward solid food but varied substantially in magnitude among individuals.

Comparison of dog-level abundance profiles across host-associated taxa revealed partial concordance among individuals. Some dogs repeatedly showed high abundance for multiple host-associated species, whereas others showed elevated abundance for only a single species or none of the focal taxa. This indicates that individualized microbiome signatures emerge through a combination of taxon-specific dynamics and overlapping host-level patterns.

Maternal background was assessed by analyzing Ag1 samples separately, using litter identity as a proxy. Litter explained 10.8% of variation in ONT 16S profiles (R² = 0.1084, p = 0.171) and 13.1% in mWGS profiles (R² = 0.1305, p = 0.062). These results suggest a modest litter-associated effect at the neonatal stage, with a stronger signal in mWGS data.

### Supplementary Results 3. Virome composition, host associations, and developmental restructuring

Viral contigs were identified from Illumina metagenomic co-assemblies using stringent filtering criteria. Of 5,109 initially detected viral contigs, 2,631 high-confidence viral contigs were retained, including 2,447 classified as prokaryotic viruses. High-confidence host predictions were available for 957 viral contigs corresponding to 191 bacterial genera.

Virome richness increased substantially across dietary stages. Using a threshold of >10 mapped reads per age group, the number of detected viral contigs increased from 807 in Ag1 to 2,434 in Ag4. Relative abundance summaries similarly indicated increasing representation of viral reads across development, with viral read fractions ranging from 16.5% to 30.2%.

Temperate phages dominated the virome in all age groups, whereas virulent phages constituted a smaller but persistent component. Viral lifestyle composition varied among predicted bacterial hosts. Bacteroides and Faecalibacterium were associated primarily with temperate phages, whereas Enterococcus and Escherichia showed more balanced associations with both temperate and virulent viruses.

Virus–host interaction networks showed heterogeneous connectivity patterns. Escherichia displayed the highest number of viral associations, whereas genera such as Paenibacillus, Shigella, and Streptococcus were associated with fewer viral taxa. Network visualization was restricted to high-confidence interactions supported by at least two viral contigs.

Viral species abundance patterns showed strong developmental structuring. Araarivirus arawn, Gemsvirus gv5004652, and Quenstivirus Q54 increased progressively with age, whereas Immutovirus immuto, Vibrio phage X29, and Gilsonvirus comrade were more abundant during breastfeeding stages. Other viral taxa, including Geobacillus virus E3 and Lactobacillus phage LfeSau, appeared predominantly during transitional or later dietary stages.

Predicted bacterial host abundances derived from viral host assignment showed broad concordance with whole-community metagenomic profiles. Later-stage anaerobic genera, including Bacteroides, Faecalibacterium, Ruminococcus, and Prevotella, increased during maturation, whereas early colonizers such as Escherichia declined across development. Correlation analyses further supported concordance between CHERRY-derived host estimates and Kraken2-derived bacterial abundance profiles for several taxa, including Escherichia albertii, Bifidobacterium animalis, Clostridium kluyveri, and Cronobacter dublinensis.

### Supplementary Results 4. Pathway-level functional restructuring across dietary transitions

KEGG ortholog profiles derived from mWGS data were used to assess functional maturation across consecutive dietary transitions. Differential KO abundance was tested with DESeq2, and significantly differentially abundant KOs (FDR < 0.05) were used for pathway-level over-representation analysis against the detected KO universe. Directionality of enriched pathways was summarized using KO-level log2 fold changes within each pathway.

The Ag1→Ag2 transition showed the most extensive pathway-level signal. Enriched pathways included flagellar assembly (fold enrichment ≈ 2.0, q ≈ 2 × 10⁻⁹), biofilm formation (≈ 2.0, q ≈ 7 × 10⁻⁹), bacterial chemotaxis (≈ 1.9, q ≈ 7 × 10⁻⁴), and two-component signal transduction systems (≈ 1.36, q ≈ 3 × 10⁻⁹). Core metabolic pathways were also enriched, including carbon metabolism (≈ 1.35, q ≈ 1 × 10⁻⁴), biosynthesis of cofactors (≈ 1.36, q ≈ 6 × 10⁻⁶), and quinone biosynthesis (≈ 1.8, q ≈ 7 × 10⁻⁴).

KO-level directionality showed that colonization-associated pathways were coherently biased toward Ag1, with median termLFC values around −2 for flagellar assembly and biofilm formation. In contrast, enriched metabolic pathways showed smaller and more heterogeneous shifts, typically with median termLFC values between −1 and 0.

The Ag2→Ag3 transition showed fewer enriched pathways, consistent with reduced functional turnover after initial dietary diversification. Enrichment was concentrated in regulatory and cell-envelope–associated processes, including two-component systems and teichoic acid biosynthesis, with KO-level shifts centered closer to zero than in the Ag1→Ag2 comparison.

The Ag3→Ag4 transition showed a restricted but coherent enrichment pattern. The phosphotransferase system was among the strongest enriched pathways (fold enrichment ≈ 3.0), although its median termLFC was modest (≈ −0.6), suggesting quantitative rebalancing rather than wholesale functional gain or loss. Nitrotoluene degradation showed stronger enrichment (≈ 4–8-fold) and a clearer Ag4-biased directionality (median termLFC ≈ +1.4), indicating targeted late-stage metabolic adaptation.

**Supplementary Results 5. Platform effects and copy-number normalization in matched ONT–mWGS profiles**

ONT full-length 16S taxonomic profiles were benchmarked against matched Illumina mWGS profiles to assess platform-specific bias and the effect of 16S rRNA gene copy-number normalization. Bray–Curtis PCoA showed clear separation by sequencing strategy, with mWGS profiles forming a distinct cluster from both raw and rrnDB-normalized ONT 16S profiles. This indicated that platform effects exceeded biological stage effects in global community structure.

Across the full dataset, 5,059 bacterial species were detected. Of these, 2,832 were detected exclusively by mWGS, 8 exclusively by ONT 16S, and 2,219 were shared between platforms. Thus, mWGS recovered a substantially broader taxonomic repertoire, whereas most 16S-detected taxa belonged to a shared core also observed in shotgun metagenomes.

PERMANOVA with sequencing method and age group as explanatory variables showed that both factors were significant at all taxonomic ranks (p = 0.001), using marginal tests with permutations restricted within host identity. The fraction of variance explained by method decreased with taxonomic resolution, from R² = 0.44 at phylum level to R² = 0.165 at species level. In contrast, age-associated variance increased from R² = 0.072 at phylum level to R² = 0.137 at species level. These results indicate that methodological effects dominate at shallow taxonomic levels, whereas biological stage effects become more comparable at genus and species resolution.

Species-level mixed-effects comparisons showed that rrnDB normalization reduced deviation from mWGS for 42 abundant species but increased deviation for 29 abundant species. Species showing reduced deviation included *Escherichia* *coli* and *Peptacetobacter* *hiranonis*. In contrast, several abundant gut-associated taxa, including *Faecalibacterium* *prausnitzii*, *Phocaeicola* *vulgatus*, *Dorea* *longicatena*, *Anaerostipes* *caccae*, and *Blautia* *parvula*, showed increased deviation after normalization.

Community-level concordance was assessed by calculating Bray–Curtis dissimilarity between each ONT V1–V9 profile and its matched mWGS profile. Non-normalized ONT profiles had a mean Bray–Curtis distance of 0.606 (IQR = 0.210), whereas rrnDB-normalized profiles had a higher mean distance of 0.712 (IQR = 0.207). This increase was statistically significant (paired Wilcoxon signed-rank test, p = 2.7 × 10⁻⁹), and most samples showed increased distance after normalization. Restricting the analysis to species with direct rrnDB entries yielded similar results, with normalized profiles showing higher dissimilarity than non-normalized profiles (mean 0.725 vs 0.623; paired Wilcoxon p = 3.4 × 10⁻⁹).

### Supplementary Results 6. Functional inference from long-read 16S profiles

Functional predictions from ONT full-length 16S profiles were benchmarked against matched mWGS-derived KEGG ortholog profiles across 142 paired samples. Three workflows were evaluated: NanoCLUST consensus sequences with default PICRUSt2 settings, NanoCLUST consensus sequences after strand correction and chimera removal with stringent placement criteria, and NanoASV-generated ASVs with the same stringent thresholds.

Read retention was defined as the fraction of total ONT reads per sample originating from ASVs or consensus sequences successfully placed and retained by PICRUSt2 after alignment and NSTI filtering. Default NanoCLUST retained 0.47–0.57 of reads across age groups, with relatively high variability. Curated NanoCLUST improved retention to 0.57–0.72. NanoASV showed the highest retention, ranging from 0.78 to 0.91, with lower variability across age groups. Relative to default NanoCLUST, optimized preprocessing increased the proportion of reads contributing to functional inference by approximately 20–35 percentage points.

KO detection accuracy was evaluated independently of abundance scaling by comparing predicted and mWGS-observed KO presence. Default NanoCLUST showed high precision (0.889–0.905) but lower recall (0.670–0.769), yielding F1 scores of 0.763–0.819. Curated NanoCLUST maintained high precision (0.874–0.904) while improving recall to 0.731–0.812, resulting in F1 scores of 0.791–0.855. NanoASV produced higher recall (0.862–0.900) but lower precision (0.729–0.791), with F1 scores of 0.787–0.837.

Inference-level concordance was assessed by correlating signed log10-transformed p-value vectors from KO-wise Wilcoxon rank-sum tests performed independently on PICRUSt2 and mWGS profiles. Across Ag1→Ag2, Ag2→Ag3, and Ag3→Ag4 contrasts, observed correlations were consistently separated from permutation-based null expectations. Spearman correlations ranged from 0.338 to 0.755 and Pearson correlations from 0.408 to 0.776 across workflows and contrasts. In Ag1→Ag2, Spearman correlations were 0.755 for default NanoCLUST, 0.737 for curated NanoCLUST, and 0.738 for NanoASV, with corresponding Pearson correlations of 0.776, 0.755, and 0.766. Permutation-based correlations were centered near zero (|ρ| < 0.01), and all observed correlations were significantly greater than null expectations (one-sided empirical p = 0.0099; 999 permutations).
