## Supplementary figures and images for "Early-life canine gut microbiome maturation follows a shared age–diet trajectory within persistent host-specific structure"

### Supplementary Figure S1

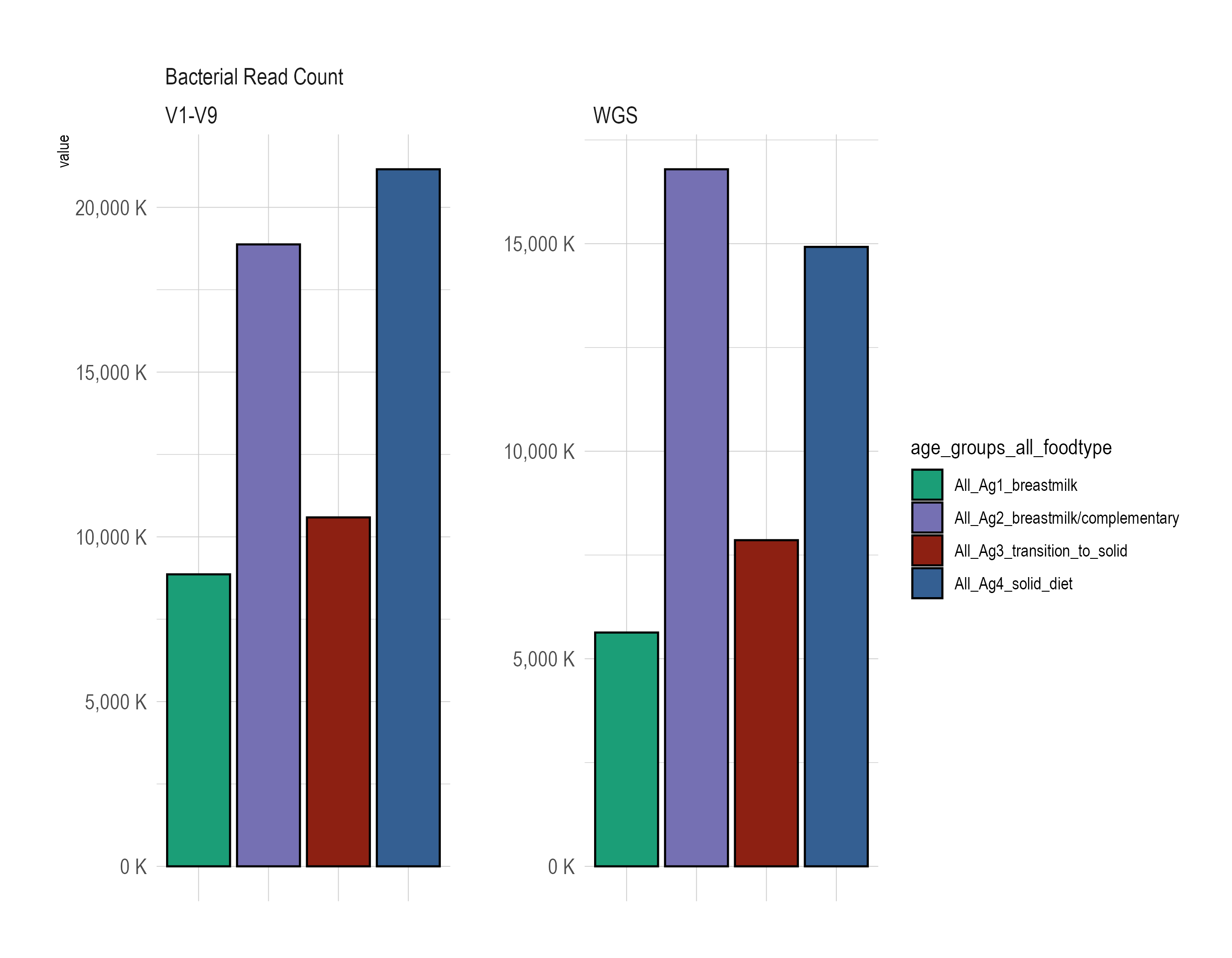

### Supplementary Figure S2

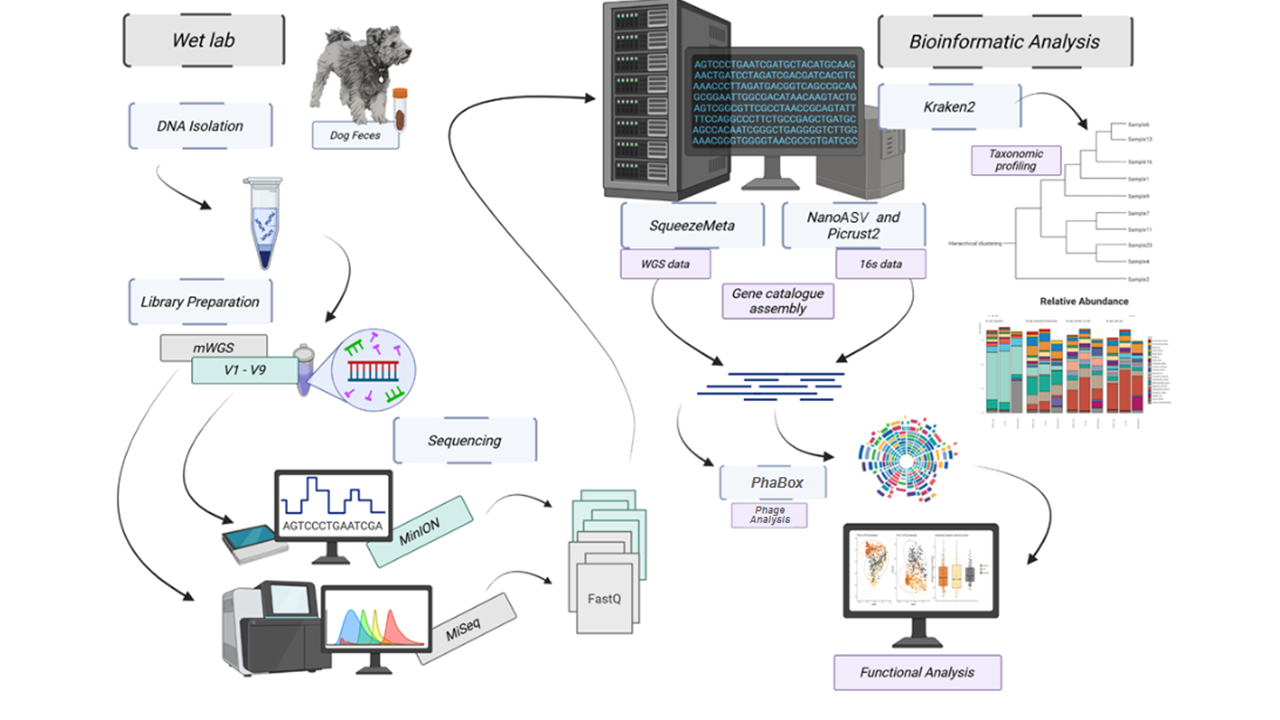

### Supplementary Figure S3

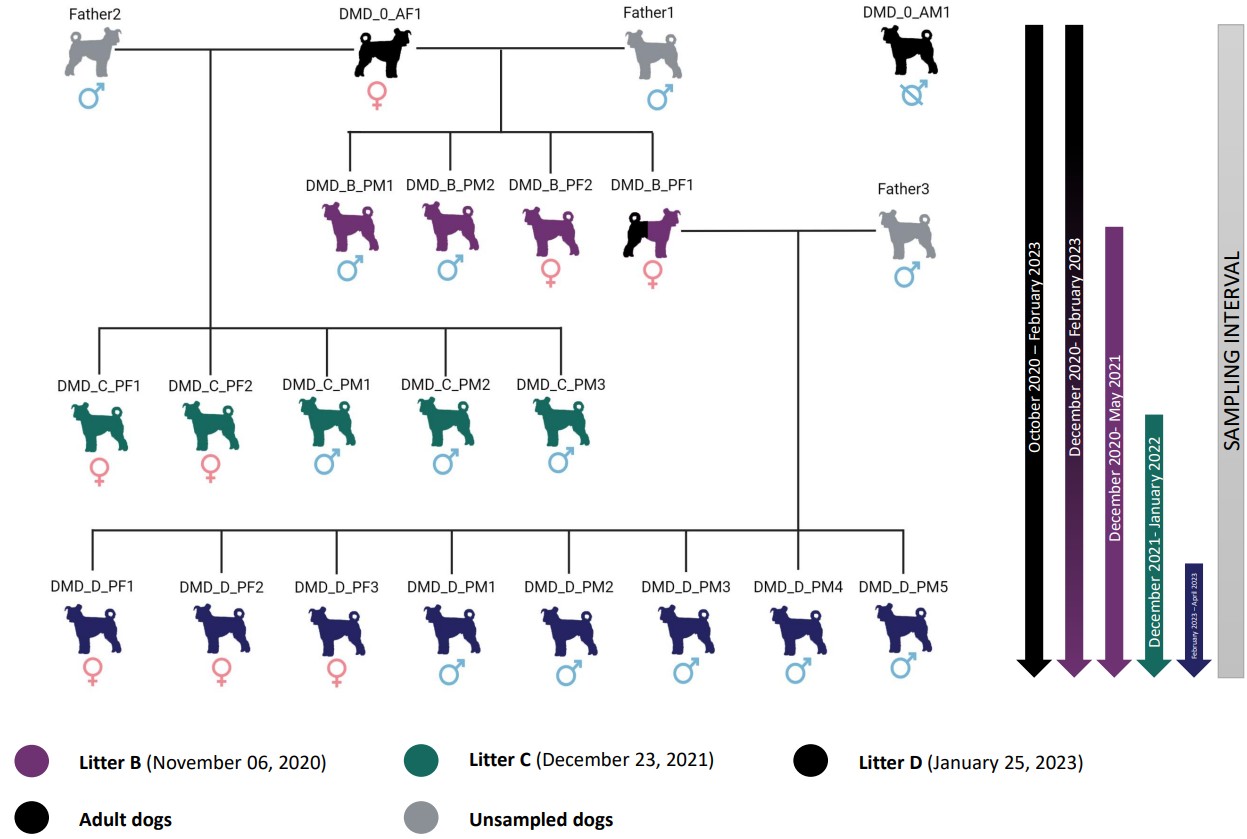

### Supplementary Figure S4

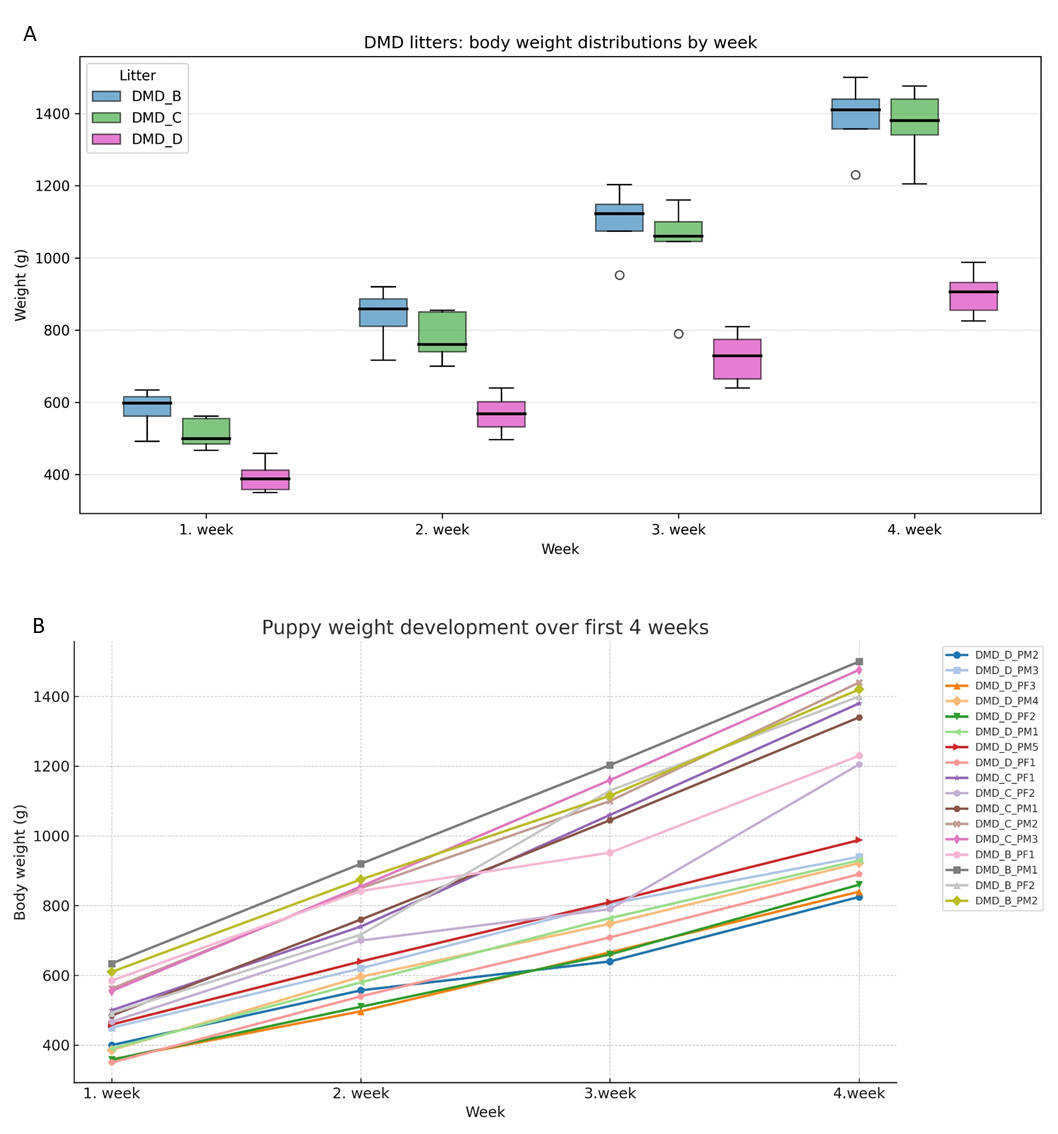
